# An atlas of the developing *Tribolium castaneum* brain reveals conserved anatomy and divergent timing to *Drosophila melanogaster*

**DOI:** 10.1101/2021.11.30.470557

**Authors:** Max S. Farnworth, Gregor Bucher, Volker Hartenstein

## Abstract

Insect brains are formed by conserved sets of neural lineages whose fibres form cohesive bundles with characteristic projection patterns. Within the brain neuropil these bundles establish a system of fascicles constituting the macrocircuitry of the brain. The overall architecture of the neuropils and the macrocircuitry appear to be conserved. However, variation is observed e.g., in size and shape and timing of development. Unfortunately, the developmental and genetic basis of this variation is poorly understood although the rise of new genetically tractable model organisms such as the red flour beetle *Tribolium castaneum* allows the possibility to gain mechanistic insights. To facilitate such work, we present an atlas of the developing brain of *T. castaneum*, covering the first larval instar, the prepupal stage and the adult, by combining wholemount immunohistochemical labelling of fibre bundles (acetylated tubulin) and neuropils (synapsin) with digital 3D reconstruction using the TrakEM2 software package. Upon comparing this anatomical dataset with the published work in *D. melanogaster*, we confirm an overall high degree of conservation. Fibre tracts and neuropil fascicles, which can be visualized by global neuronal antibodies like anti-acetylated tubulin in all invertebrate brains, create a rich anatomical framework to which individual neurons or other regions of interest can be referred to. The framework of a largely conserved pattern allowed us to describe differences between the two species with respect to parameters such as timing of neuron proliferation and maturation. These features likely reflect adaptive changes in developmental timing that govern the change from larval to adult brain.

## Introduction

The evolutionary history of animals is reflected in the structure of their brains, which process sensory inputs and mediate adaptive behaviours (Sterling & Laughlin, 2015). Brain anatomy therefore constitutes an important aspect of animal evolution. Following this rationale, previous studies of insect brains have related differences in the volume and structure of various brain domains to specific features in the animals' environment, sensory capabilities, and the resulting behavioural repertoires (Farris, 2013; Montgomery et al., 2021; Pisokas et al., 2020). Most of such studies have focused on four “classic” brain domains, or neuropils, i.e., antennal lobes (AL), mushroom bodies (MB), optic lobes (OL) and central complex (CX), which in most species can be visible even without specific labels (Dujardin, 1850; Strausfeld, 2012). This approach has generated a wealth of data that was used to relate a brain domain’s size to its relevance in function and predict patterns of adaptation and clade shifts during insect evolution.

Despite their crucial role in brain function, less attention has been dedicated to brain macro-connectivity, i.e., the systems of nerve fibre bundles that innervate and interconnect specific brain domains. Documenting and interpreting fibre tracts in the context of animal development and evolution has a long tradition in vertebrate neuroanatomy, where all structurally defined domains (e.g., brainstem nuclei, cortical layers and areas) are bounded and/or interconnected by specific axon tracts. Fibre systems that make up the macro-connectivity of the vertebrate brain, such as the cortico-thalamic tract or spino-cerebellar tract, are prominent research targets, from comparative neuroanatomy to clinical research (Leyva-Díaz & López-Bendito, 2013).

Comparing the pattern of fibre connections in insect brains would be an essential addition because tracts not only represent the structural units of macro-connections within the brain, but also reflect the developmental-genetic history of the neurons by which they are formed. Indeed, the insect brain is formed by neural lineages, which comprise clusters of neurons derived by an invariant developmental pattern from one neural stem cell (neuroblast) each. Neurons of a given lineage, or parts thereof (hemilineage, sublineage), form bundles that make up the systems of tracts and fascicles of the brain (Figure 1, Lee, 2017; Spindler & Hartenstein, 2010).

**Figure 1:**
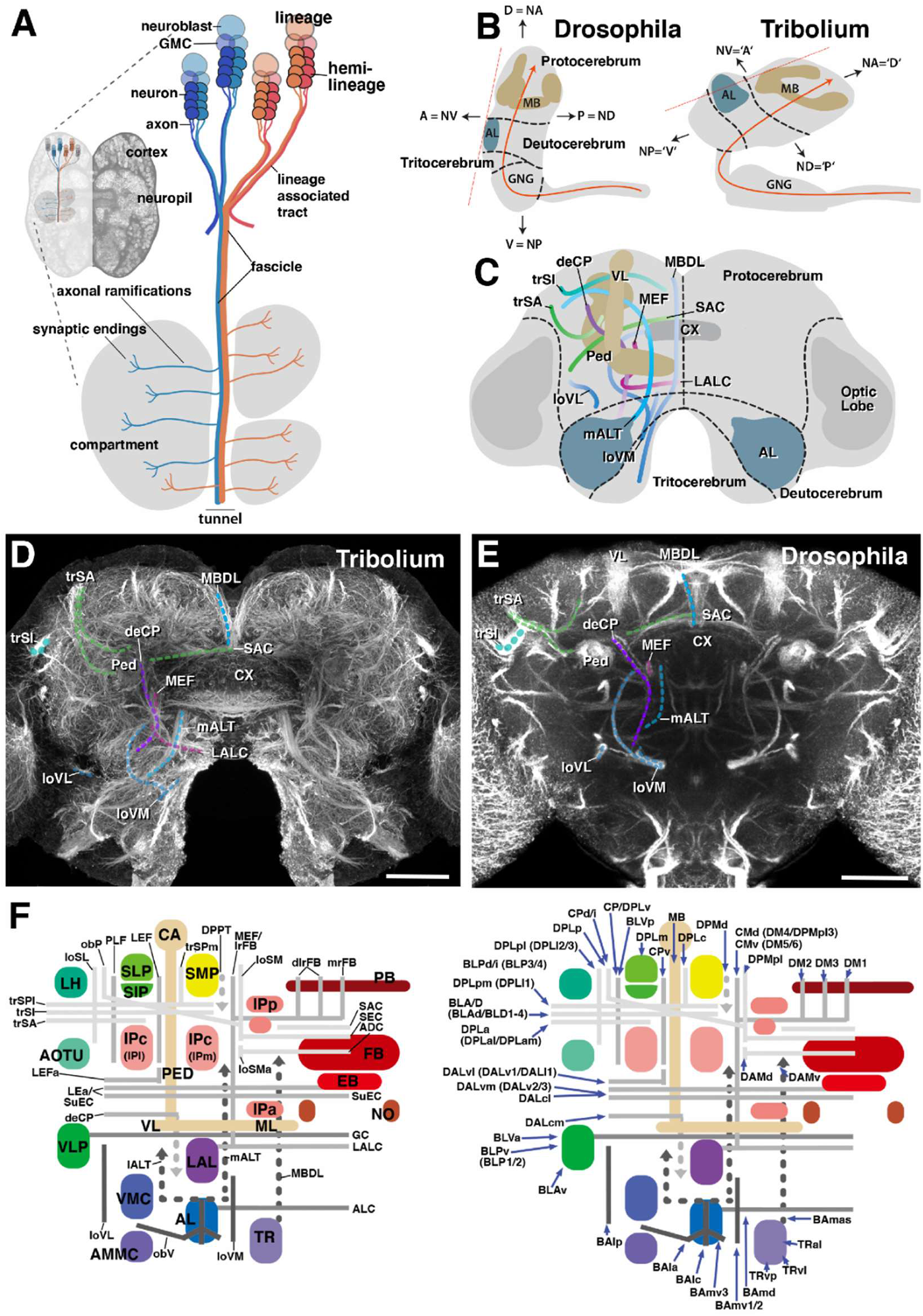
Fundamentals of insect brain anatomy. (A) Schematic of larval brain illustrating the features of neural lineage formation [lineage, hemilineage, neuroblast, ganglion mother cell (GMC), neuron, lineage associated tract] and the spatial relationship between lineage associated tracts and cortex, neuropil and neuropil compartments. (B) Schematics of brains of *D. melanogaster* and *T. castaneum* (lateral view), showing neuraxis (orange line), arrangement of brain neuromeres [protocerebrum, deutocerebrum, tritocerebrum, gnathal ganglia (GNG)] and angle of frontal plane (orange dashed line) relative to neuraxis. (C) Schematic of insect brain (anterior view, shape of adult *T. castaneum* brain) with neuromeres and assemblage of representative neuropil fascicles in different colors. (D, E) Z-projections of frontal confocal section of adult *T. castaneum* (D) and *D. melanogaster* brain (E) labeled with global neuronal marker (*T. castaneum*: Tubulin; *D. melanogaster*: Neuroglian). Segments of homologous fascicles shown in the schematic (C) are shaded and annotated. (F) Schematic *T. castaneum* brain representations of fascicles and compartments (left) as well as tracts (right) now described and implemented in this study. Colours as in the models. For abbreviations see Table 2. Scale bars: 25µm (D, E).

The conservation of insect neuroblasts and their lineages has been studied intensely (Bate, 1976; Biffar & Stollewerk, 2014; Boyan & Reichert, 2011; Broadus & Doe, 1995; Lovick et al., 2017; Thomas et al., 1984; Truman & Ball, 1998; Urbach & Technau, 2003a). Differences in brain anatomy between taxa can be related to modifications of parameters of neuroblast patterning and proliferation. These include the duration of the proliferative period (Truman & Ball, 1998), the multiplication of discrete neuroblasts (e.g. 500 MB neuroblasts in *Apis mellifera* versus four in *D. melanogaster*, Farris, 2013; and two in *T. castaneum*, Farris & Roberts, 2005; Kebschull et al., 2020; Strausfeld, 2005, this work), as well as lineage degeneration by programmed cell death (Pop et al., 2020; Prieto-Godino et al., 2020). Insect brains, therefore, offer the opportunity to examine the developmental basis of brain evolution in the framework of conversed developmental units and adult anatomy.

A detailed representation of the lineages and their associated tracts has been generated for the developing *D. melanogaster* brain. Individual lineages were mapped by clonal labelling techniques (Larsen et al., 2009; Lee et al., 2020; Lovick et al., 2017; Pereanu & Hartenstein, 2006; Wong et al., 2013; Yu et al., 2013), and the pattern of lineage associated tracts was reconstructed from early larval into the adult stage (Hartenstein et al., 2015; Lovick et al., 2013; Pereanu et al., 2010; Pereanu & Hartenstein, 2006). In conjunction with markers that highlight synapses, a system of neuropil compartments, fascicles, and lineage-associated tracts has been defined (Figure 1). This description of the *D. melanogaster* brain has been the basis for functional studies (e.g. Hardcastle et al., 2021; Omoto et al., 2018) but at the same time is the basis for comparative developmental studies.

In terms of functional genetic tools, the red flour beetle, *Tribolium castaneum*, is the second-most advanced insect model, as techniques such as CRISPR, GAL4/UAS, systemic and parental RNAi and large-scale screening opportunities are well established (Berghammer et al., 1999; Brown et al., 2009; Bucher et al., 2002; Gilles et al., 2015; Schinko et al., 2009; Schinko et al., 2010; Schmitt-Engel et al., 2015). In recent years, *T. castaneum* has been used for neurobiological research, examining the heterochronic development and evolution of the central complex (Farnworth et al., 2020; Garcia-Perez et al., 2021; He et al., 2019) and neuroblast patterning (Biffar & Stollewerk, 2014, 2015). Unfortunately, brain compartments, fascicles and tracts have not been comprehensively described, limiting the breadth of comparison with other species. Standard neuropils have been mapped in *T. castaneum* (Dreyer et al., 2010; Koniszewski et al., 2016) and, in another Coleopteran, the dung beetle, the compartments and major fascicles of the adult brain have been determined, which in part may be generalized along Coleopterans (Immonen et al., 2017). In addition, there is a database documenting the volumes of the four brain neuropils in adult Coleopterans (Kollmann et al., 2016). However, very little data exist on the neuropils and microcircuitry of Coleopteran developmental stages (see Farnworth et al., 2020 and Wegerhoff & Breidbach, 1992 for details on the larval central complex in Coleoptera, as well as Koniszewski et al., 2016 for a general map of the first larval stage brain in *T. castaneum*).

A more detailed and tract-oriented brain anatomy in *T. castaneum* will provide a framework that will greatly enhance neurobiological and developmental studies in this species. We were particularly interested to ascertain whether the suggested conservation of the basic pattern of lineages, posited almost 40 years ago (Thomas et al., 1984), is reflected when comparing the tract anatomy of *T. castaneum* and *D. melanogaster*. Importantly, we wanted to detect and define differences thereby providing models for studying the developmental basis of brain evolution. To reach this aim, we have used anti-acetylated tubulin and anti-synapsin antibodies in conjunction with several other markers to reconstruct brain anatomy. Acetylated tubulin labels neuronal somata and most parts of the neurites (not growth cones, Perdiz et al., 2011; Robson & Burgoyne, 1989), while Synapsin stainings revealed synapse-rich regions of the brain. We generated an atlas of compartments and fascicles at the first larval, prepupal and adult stage, identified putative connections of tracts with lineages at the larval stage, and we drew a comparison to the *D. melanogaster* atlas. We found a strong conservation between the two species, with respect to compartment and fascicle anatomy down to the pattern of tracts. We describe each element in detail, with an emphasis on the larval stage where the pattern of tracts is most clearly visible. Importantly, we found considerable differences with respect to the spatio-temporal pattern of neuron proliferation and differentiation. We discuss the differences in the context of evolution of sense organ development and holometaboly.

## Material and Methods

### Data availability and deposition

We deposited all original data on which the figures are based on, specific annotated stacks as well as models of the compartments of the L1 and adult brain on www.insectbraindb.org (Heinze et al., 2021). Upon technical expansion of the 3D viewers and interactive experiments, we will include tracts and fascicles into the 3D models. At the moment, all 3D models can be accessed using Fiji and guidelines highlighted in the original TrakEM2 publication (Cardona et al., 2012).

Each time point and staining type (synapsin or tubulin staining) is represented by at least N=3 individuals.

### Stocks, handling and staging

*D. melanogaster:* All flies used were OregonR wildtype. Stock keeping and staging followed established procedures (Sullivan et al., 2000).

*T. castaneum:* We used a transgenic line that was generated by piggyBac transposition to insert a 6XP3-dsRedExpress construct, as 6XP3 drives expression in eye tissue as well as in most glial cells (Koniszewski et al., 2016).

Animals were kept either at 28°C or 32°C for egg lays under standard conditions (Brown et al., 2009). For the L1 stage, we collected embryos and put them onto 300 mm gaze, through which hatched larvae will crawl through. We collected L1 with a maximum age of 1 h post-hatching, as to ensure homogeneity in staging.

The prepupal stage was determined as in Farnworth et al. (2020), i.e., by its highly characteristic crescent-shaped body position and reduced body mobility. As this stage only covers a few hours, we saw this as the most exact stage in larval development besides L1.

We only took adult beetles that hatched less than ~12 hours ago, as identified by their light-blown/yellow, translucent cuticle.

### Dissections

In essence, we dissected brains using Dumont No. 5 forceps for a maximum of 30 minutes, storing them in ice-cold phosphate-buffered saline (PBS) until fixations started.

*D. melanogaster* brain dissections followed standard procedures (Lovick et al., 2017; Sullivan et al., 2000).

*T. castaneum* larval L1 brain dissections largely followed procedures described in Hunnekuhl et al. (2020). The larval brain is located between the posterior end of the head capsule and the first thoracic segment (see Figure 1). With that in mind, L1 brains were dissected by removing the abdomen, removing the anterior head structures, and pushing out the brain including most ganglia by pressing on the cuticle. Prepupal brains were dissected very similarly, but instead of pushing out the brain, we freed the brain by removing the remaining body tissue bit by bit. Adult brains were dissected by, first, removing the whole head capsule. Then, forceps were inserted into the hole as ventral as possible. The cuticle was cut towards anterior up to and including the eyes on both sides and lifted from the remaining head capsule. Then, the head was turned slightly on its top, such that the mouth parts were directed to the experimenter. Each half of the dorsal side of the head capsule was carefully removed, leaving brain tissue embedded in a lot of fat and tracheal tissue. This was carefully removed to liberate the adult brain.

### Fixation, Immunohistochemistry and Mounting

*T. castaneum brain stainings*: All brains for tubulin staining were fixed in self-made Zinc-Formaldehyde (ZnFA) solution (as in Ott, 2008) while we used a freshly prepared solution of 4% Methanol-free Formaldehyde (FA; #28908 ThermoFisher Scientific) diluted in 1x PBS for synapsin staining. ZnFA was used as we observed a loss of microanatomy with respect to fibre and tract anatomy when using FA in tubulin staining, which, however, resulted in better penetration. L1 brains were fixed for 1 h, prepupal brains for 1 h 15 min, and adult brains for 1 h 30 min, all with ZnFA at room temperature, and with FA at 4°C with slight agitation on an orbital shaker. Fixations and all washings without detergent were done in black glass wares or 9-well PYREX™ Spot Plates (ThermoFisher Scientific, MA, USA) to avoid sticking brains onto plastic.

Fixation was followed by three rinses and three 15-minute washes in ice-cold PBS (all washing and agitation steps occurred on an orbital shaker). Then brains fixed in FA were dehydrated with ice-cold Ethanol/PBS mixtures in 12.5% steps and brains fixed in ZnFA were dehydrated as described in Ott (2008). During dehydration all brains were transferred to 0.5 ml Eppendorf tubes and stored over night at −20°C. Rehydration occurred in 12.5% steps again or following Ott (2008) with transfer to glass ware again, followed by 3 rinses in ice-cold PBS and then several three 15-minute washes in PBS-Triton-X-100 (0.1% for L1, 0.3% for prepupa and adults) or PBS-DMSO (1 %) for ZnFA-fixed brains. This was followed by a 3 h blocking step with 5 % NGS (G9023, Merck, Darmstadt Germany) in the respective washing buffer, at room temperature with agitation. The primary antibodies diluted in 2 % NGS in wash buffer were incubated at 4°C with agitation at following durations and concentrations (see Table 1 for antibody details): The antibody targeting synapsin was used on L1 brains at 1:25 dilution for 24 h, and at 1:20 dilution for 72 h on prepupal and adult brains. The acetylated Tubulin antibody was used at a 1:50 dilution for 24 h in L1 brains, 1:100 dilution for 5 days in prepupal and adult brains. The RFP antibody against DsRed-Express was used in a 1:500 dilution combined with synapsin or tubulin at the same times in the respective stages. Antibody incubation was followed by three rinses and three 15 min washes in the respective wash buffers. Secondary antibodies were incubated in 1% NGS in wash buffers at 4°C with agitation with a 1:400 dilution at the same durations as the first antibodies (goat anti-mouse-Alexafluor-488, A-11017; goat anti-rabbit-Alexafluor-555, A-21430; ThermoFisher Scientific/Invitrogen, MA, USA). Then, the brains were rinsed three times, incubated with DAPI (D1306, ThermoFisher Scientific, MA, USA) at 1:1000 (prepupa, adult) or 1:1500 dilution (L1) for 30 minutes at room temperature, followed by one rinse and four 15 minute washes. This was lastly followed by a PBS wash for 2 h (prepupa and adult) or 1 h (L1). The brains were then mounted on frosted and 90° ground object slides (AGAA000001##12E, ThermoFisher Scientific, MA, USA) inside paper hole reinforcements (3510, Avery Zweckform, Oberlaindern, Germany; 2 for prepupa and adult, 1 for L1, to avoid compression) with 25×40 mm coverslips of #1.5 thickness. Brains were cleared using RapiClear 1.47 medium (#RC147001, SUNJin Lab, Hsinchu City, Taiwan) by applying it for 2 mins followed by normal mounting. Following a 2 h incubation at 4°C they were then frozen at −20°C for further use.

**Table 1:** Antibodies used.

| Antibody | Immunogen | Manufacturer, Cat#, RRID, Species, Clonality | Dilution |
| --- | --- | --- | --- |
| acetylated tubulin | acetylated tubulin from the outer arm of *Strongylocentrotus purpuratus* (sea urchin) | Sigma-Aldrich, T7451, AB_609894, mouse, monoclonal | 1:50-100 |
| Bruchpilot | Head homogenate | DSHB, nc82, AB_2314866, mouse, monoclonal | 1:10 |
| DN-Cadherin | Recombinant fusion protein with extracellular region of N-cadherin from cadherin repeats 2 - 8 | DSHB, DN-Ex #8, AB_528121, rat, monoclonal | 1:10 |
| Neuroglian | Membrane proteins of cultured embryonic *D. melanogaster* neurons | DSHB, BP 104, AB_528402, mouse, monoclonal | 1:10 |
| Neurotactin | Membrane proteins of cultured embryonic *D. melanogaster* neurons | DSHB, BP 106, AB_528404, mouse, monoclonal | 1:10 |
| RFP | Recombinant full length protein corresponding to RFP | Abcam, ab62341, AB_945213, rabbit, polyclonal | 1:500 |
| synapsin | *D. melanogaster* Synapsin glutathione-S-transferase fusion protein | E Buchner University of Wuerzburg, 3C11 (SYNORF1), RRID: AB_528479, mouse, monoclonal | 1:20-25 |

**Table 2:**
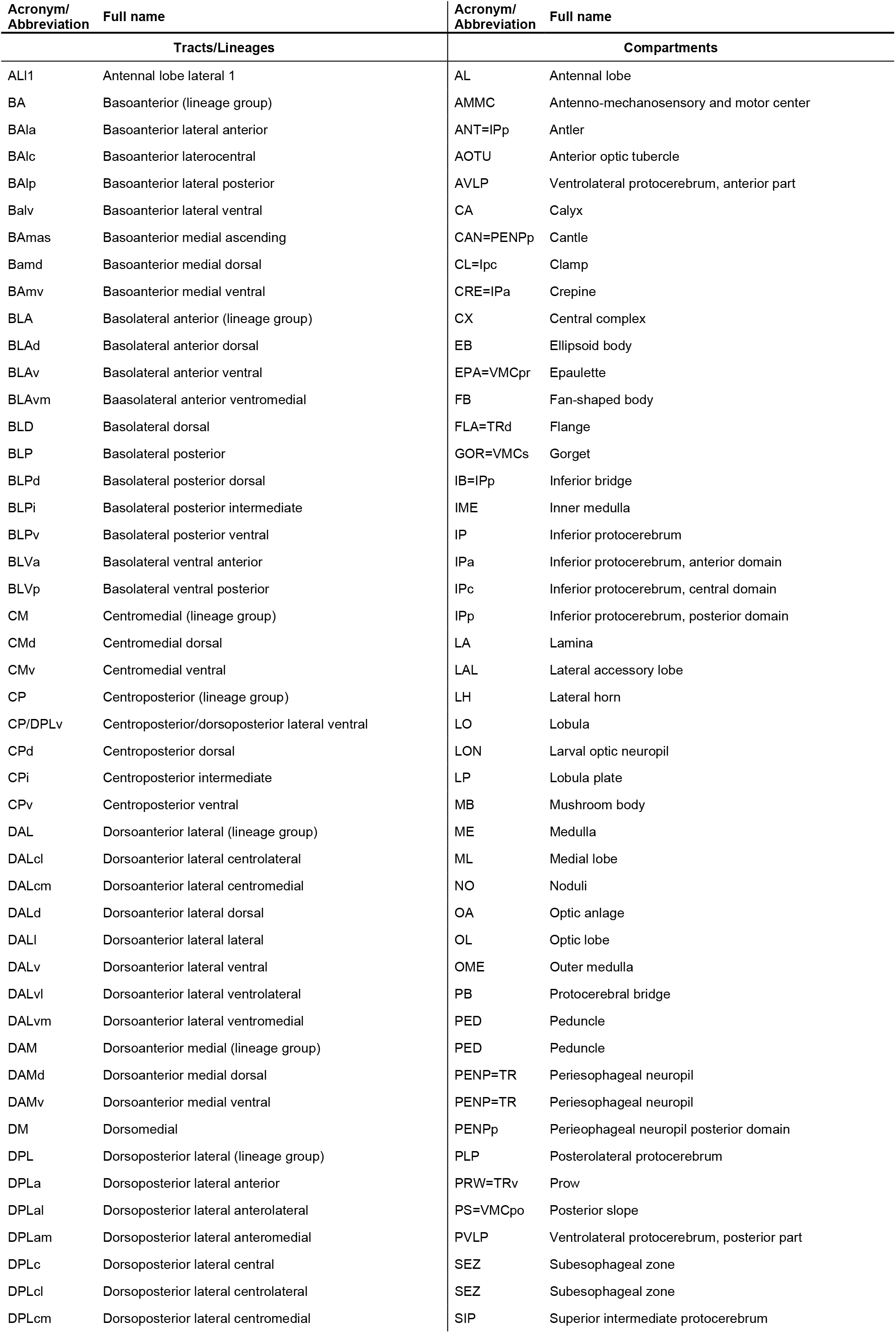
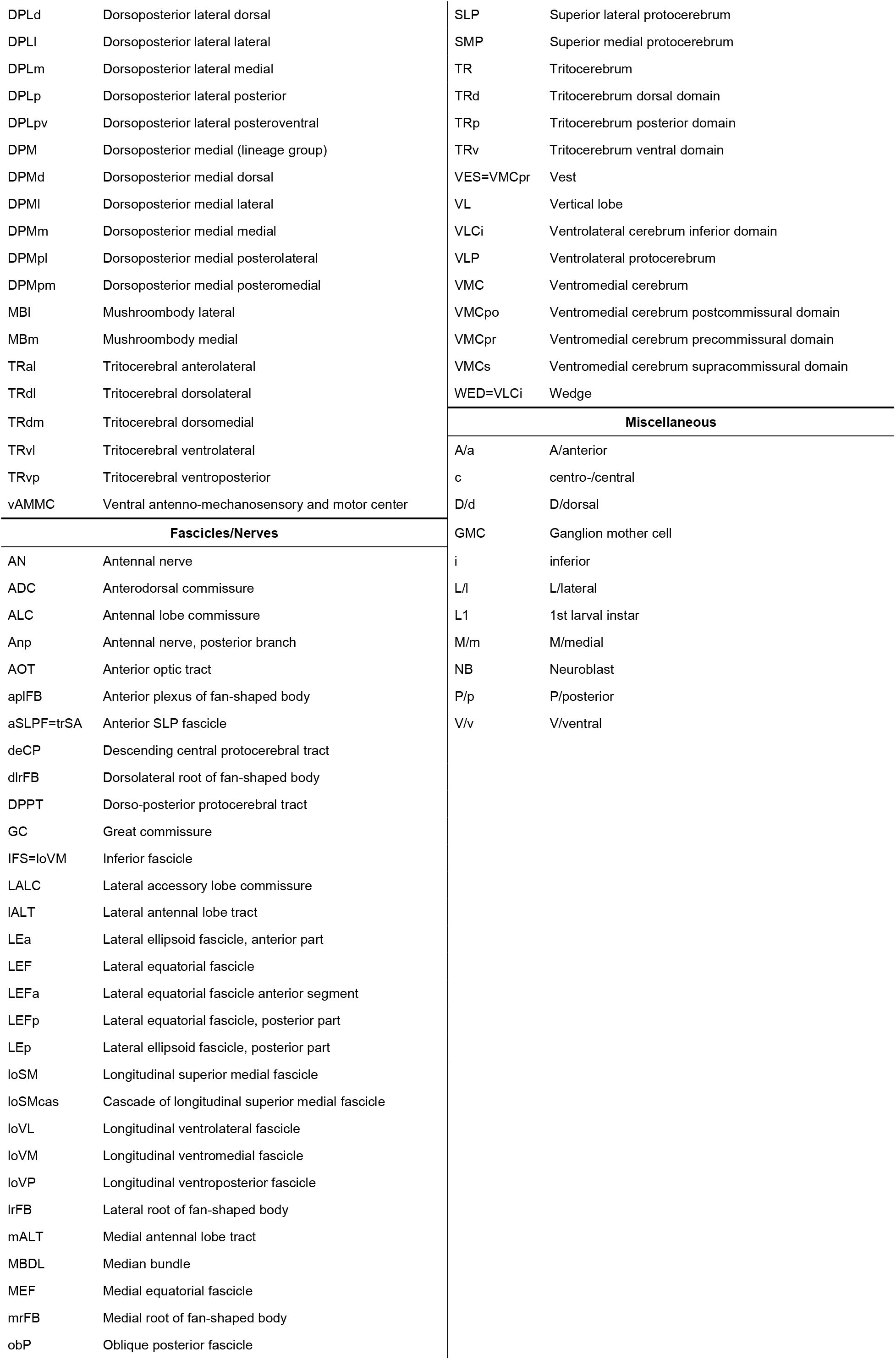
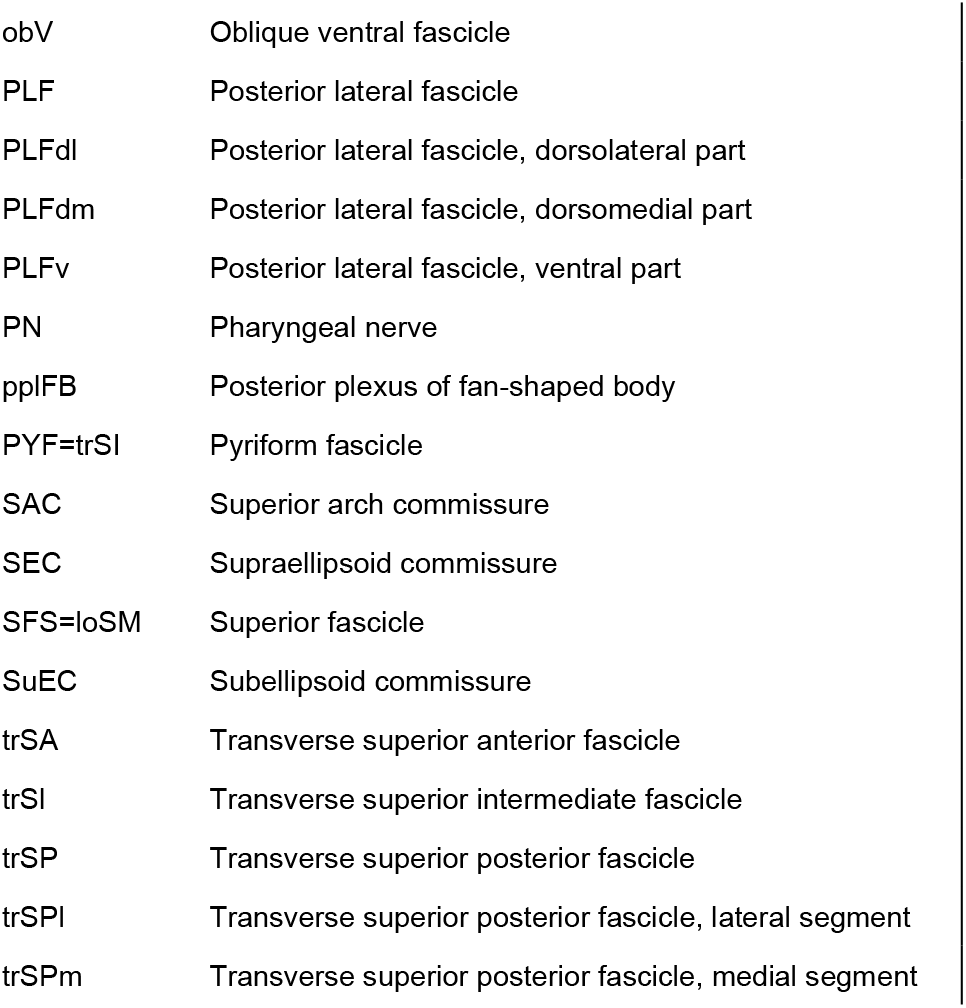
Acronyms and abbreviations used.

| Acronym/<br>Abbreviation | Full name | Acronym/<br>Abbreviation | Full name |
| --- | --- | --- | --- |
| Tracts/Lineages |  | Compartments |  |
| ALI1 | Antennal lobe lateral 1 | AL | Antennal lobe |
| BA | Basoanterior (lineage group) | AMMC | Antenno-mechanosensory and motor center |
| BAla | Basoanterior lateral anterior | ANT=IPp | Antler |
| BAlc | Basoanterior laterocentral | AOTU | Anterior optic tubercle |
| BAlp | Basoanterior lateral posterior | AVLP | Ventrolateral protocerebrum, anterior part |
| Balv | Basoanterior lateral ventral | CA | Calyx |
| BAmas | Basoanterior medial ascending | CAN=PENPp | Cantle |
| Bamd | Basoanterior medial dorsal | CL=Ipc | Clamp |
| BAmv | Basoanterior medial ventral | CRE=IPa | Crepine |
| BLA | Basolateral anterior (lineage group) | CX | Central complex |
| BLAd | Basolateral anterior dorsal | EB | Ellipsoid body |
| BLAv | Basolateral anterior ventral | EPA=VMCpr | Epaulette |
| BLAvm | Basolateral anterior ventromedial | FB | Fan-shaped body |
| BLD | Basolateral dorsal | FLA=TRd | Flange |
| BLP | Basolateral posterior | GOR=VMCs | Gorget |
| BLPd | Basolateral posterior dorsal | IB=IPp | Inferior bridge |
| BLPi | Basolateral posterior intermediate | IME | Inner medulla |
| BLPv | Basolateral posterior ventral | IP | Inferior protocerebrum |
| BLVa | Basolateral ventral anterior | IPa | Inferior protocerebrum, anterior domain |
| BLVp | Basolateral ventral posterior | IPc | Inferior protocerebrum, central domain |
| CM | Centromedial (lineage group) | IPp | Inferior protocerebrum, posterior domain |
| CMd | Centromedial dorsal | LA | Lamina |
| CMv | Centromedial ventral | LAL | Lateral accessory lobe |
| CP | Centroposterior (lineage group) | LH | Lateral horn |
| CP/DPLv | Centroposterior/dorsoposterior lateral ventral | LO | Lobula |
| CPd | Centroposterior dorsal | LON | Larval optic neuropil |
| CPi | Centroposterior intermediate | LP | Lobula plate |
| CPv | Centroposterior ventral | MB | Mushroom body |
| DAL | Dorsoanterior lateral (lineage group) | ME | Medulla |
| DALcl | Dorsoanterior lateral centrolateral | ML | Medial lobe |
| DALcm | Dorsoanterior lateral centromedial | NO | Noduli |
| DALd | Dorsoanterior lateral dorsal | OA | Optic anlage |
| DALI | Dorsoanterior lateral lateral | OL | Optic lobe |
| DALv | Dorsoanterior lateral ventral | OME | Outer medulla |
| DALvl | Dorsoanterior lateral ventrolateral | PB | Protocerebral bridge |
| DALvm | Dorsoanterior lateral ventromedial | PED | Peduncle |
| DAM | Dorsoanterior medial (lineage group) | PED | Peduncle |
| DAMd | Dorsoanterior medial dorsal | PENP=TR | Periesophageal neuropil |
| DAMv | Dorsoanterior medial ventral | PENP=TR | Periesophageal neuropil |
| DM | Dorsomedial | PENPp | Perieophageal neuropil posterior domain |
| DPL | Dorsoposterior lateral (lineage group) | PLP | Posterolateral protocerebrum |
| DPLa | Dorsoposterior lateral anterior | PRW=TRv | Prow |
| DPLal | Dorsoposterior lateral anterolateral | PS=VMCpo | Posterior slope |
| DPLam | Dorsoposterior lateral anteromedial | PVLP | Ventrolateral protocerebrum, posterior part |
| DPLc | Dorsoposterior lateral central | SEZ | Subesophageal zone |
| DPLcl | Dorsoposterior lateral centrolateral | SEZ | Subesophageal zone |
| DPLcm | Dorsoposterior lateral centromedial | SIP | Superior intermediate protocerebrum |
| DPLd | Dorsoposterior lateral dorsal | SLP | Superior lateral protocerebrum |
| DPLl | Dorsoposterior lateral lateral | SMP | Superior medial protocerebrum |
| DPLm | Dorsoposterior lateral medial | TR | Tritocerebrum |
| DPLp | Dorsoposterior lateral posterior | TRd | Tritocerebrum dorsal domain |
| DPLpv | Dorsoposterior lateral posteroventral | TRp | Tritocerebrum posterior domain |
| DPM | Dorsoposterior medial (lineage group) | TRv | Tritocerebrum ventral domain |
| DPMd | Dorsoposterior medial dorsal | VES=VMCpr | Vest |
| DPMl | Dorsoposterior medial lateral | VL | Vertical lobe |
| DPMm | Dorsoposterior medial medial | VLCi | Ventrolateral cerebrum inferior domain |
| DPMpl | Dorsoposterior medial posterolateral | VLP | Ventrolateral protocerebrum |
| DPMpm | Dorsoposterior medial posteromedial | VMC | Ventromedial cerebrum |
| MBI | Mushroombody lateral | VMCpo | Ventromedial cerebrum postcommissural domain |
| MBm | Mushroombody medial | VMCpr | Ventromedial cerebrum precommissural domain |
| TRal | Tritocerebral anterolateral | VMCs | Ventromedial cerebrum supracommissural domain |
| TRdl | Tritocerebral dorsolateral | WED=VLCi | Wedge |
| TRdm | Tritocerebral dorsomedial | **Miscellaneous** |  |
| TRvl | Tritocerebral ventrolateral | A/a | A/anterior |
| TRvp | Tritocerebral ventroposterior | c | centro-/central |
| vAMMC | Ventral antenno-mechanosensory and motor center | D/d | D/dorsal |
| **Fascicles/Nerves** |  | GMC | Ganglion mother cell |
| AN | Antennal nerve | i | inferior |
| ADC | Anterodorsal commissure | L/l | L/lateral |
| ALC | Antennal lobe commissure | L1 | 1st larval instar |
| Anp | Antennal nerve, posterior branch | M/m | M/medial |
| AOT | Anterior optic tract | NB | Neuroblast |
| aplFB | Anterior plexus of fan-shaped body | P/p | P/posterior |
| aSLPF=trSA | Anterior SLP fascicle | V/v | V/ventral |
| deCP | Descending central protocerebral tract |  |  |
| dlrFB | Dorsolateral root of fan-shaped body |  |  |
| DPPT | Dorso-posterior protocerebral tract |  |  |
| GC | Great commissure |  |  |
| IFS=loVM | Inferior fascicle |  |  |
| LALC | Lateral accessory lobe commissure |  |  |
| IALT | Lateral antennal lobe tract |  |  |
| LEa | Lateral ellipsoid fascicle, anterior part |  |  |
| LEF | Lateral equatorial fascicle |  |  |
| LEFa | Lateral equatorial fascicle anterior segment |  |  |
| LEFp | Lateral equatorial fascicle, posterior part |  |  |
| LEp | Lateral ellipsoid fascicle, posterior part |  |  |
| loSM | Longitudinal superior medial fascicle |  |  |
| loSMcas | Cascade of longitudinal superior medial fascicle |  |  |
| loVL | Longitudinal ventrolateral fascicle |  |  |
| loVM | Longitudinal ventromedial fascicle |  |  |
| loVP | Longitudinal ventroposterior fascicle |  |  |
| lrfB | Lateral root of fan-shaped body |  |  |
| mALT | Medial antennal lobe tract |  |  |
| MBDL | Median bundle |  |  |
| MEF | Medial equatorial fascicle |  |  |
| mrFB | Medial root of fan-shaped body |  |  |
| obP | Oblique posterior fascicle |  |  |

|  |  |
| --- | --- |
| obV | Oblique ventral fascicle |
| PLF | Posterior lateral fascicle |
| PLFdl | Posterior lateral fascicle, dorsolateral part |
| PLFdm | Posterior lateral fascicle, dorsomedial part |
| PLFv | Posterior lateral fascicle, ventral part |
| PN | Pharyngeal nerve |
| pplFB | Posterior plexus of fan-shaped body |
| PYF=trSI | Pyriform fascicle |
| SAC | Superior arch commissure |
| SEC | Supraellipsoid commissure |
| SFS=loSM | Superior fascicle |
| SuEC | Subellipsoid commissure |
| trSA | Transverse superior anterior fascicle |
| trSI | Transverse superior intermediate fascicle |
| trSP | Transverse superior posterior fascicle |
| trSPI | Transverse superior posterior fascicle, lateral segment |
| trSPm | Transverse superior posterior fascicle, medial segment |

*D. melanogaster brain stainings*: Stainings were performed as described by Hartenstein et al. (2018). Shortly, dissected brains were fixed in a 4% paraformaldehyde solution diluted in PBS pH=7.4 for 30 min. They were then washed in a 0.1% Triton-X-100 solution diluted in 1X PBS pH=7.4 (wash buffer) for three times, each for 10 minutes. Brains were then incubated in blocking buffer (2% bovine serum albuminum (BSA) in 1X PBS pH 7.4) for 1 h at room temperature, followed by an overnight incubation at 4°C in the respective primary antibodies diluted in blocking buffer (Table 1). This was followed by three 15 min wash steps in wash buffer and a 20 min incubation in blocking buffer. Brains were incubated in secondary antibodies over night at 4°C [Alexa Fluor 546 goat anti-Mouse (#A11030; Invitrogen, Carlsbad, CA)], used at 1:500 for Bruchpilot, Neuroglian and Neurotactin, and Cy5 goat anti-Rat (#112–175-143; Jackson Immunoresearch, West Grove,PA), used at 1:400 for DN-Cadherin). Samples were again washed three times for 15 min in wash buffer and mounted using Vectashield (Vector Laboratories, Burlingame, CA).

### Antibody characterisation

The acetylated Tubulin antibody (T7451, MERCK/Sigma-Aldrich, Darmstadt, Germany) is a widely used antibody that detects acetylated α-tubulins in several distantly related species (Table 1, see Supplier information) with an antigen that is a highly conserved epitope on the α3 isoform of *Chlamydomonas* axonemal α-tubulin. Its specificity was verified by the supplier. We used it as a general structural marker to label the tract system of the *T. castaneum* brain, without making functional inferences about acetylated tubulin itself.

The synapsin antibody is a standard antibody in insect neuroanatomy that targets synapsin, a highly expressed vesicle protein at the pre-synapse. The immunogen is a fusion protein made of glutathione S‐transferase and parts of the *D. melanogaster* Synapsin (SYNORF1, Klagges et al., 1996). It has been used in many insect species, including *T. castaneum* (Dreyer et al., 2010) and other beetles (Immonen et al., 2017; Kollmann et al., 2016), to determine neuropil shape and size as well as general brain anatomy. Its specificity was previously demonstrated for *D. melanogaster* (Godenschwege et al., 2004; Klagges et al., 1996) and the staining in *T. castaneum* reflects that pattern. The epitope LFGGMEVCGL (Hofbauer et al., 2009) in a *T. castaneum* genome BLAST returns the *synapsin* gene TC002264.

The RFP antibody was used to detect dsRedExpress which was transgenically expressed in glial cells of *T. castaneum* allowing for an additional anatomical label. The RFP antibody (Abcam, ab62341) is universally used to detect RFP and derivatives, and was shown by the manufacturer to recognize recombinant RFP.

The Bruchpilot antibody, a synapse label alternative to synapsin was raised against adult *D. melanogaster* head homogenates and verified by Western Blotting (Wagh et al., 2006). Two bands were detected, belonging to the same transcript of the *bruchpilot* gene.

The DN-cadherin antibody, an overall marker for neuropil, was generated and validated for its specificity by Iwai et al. (1997). First, two major Western Blot bands were detected after DN-Cadherin protein transfection. Second, a staining was hardly detectable in homozygous *D. melanogaster* DN-cadherin mutants, but in those mutants that expressed a DN-Cadherin transgene.

The Neuroglian antibody labels secondary neurons in the adult *D. melanogaster* brain. Its specificity was verified by immunoaffinity chromatography and subsequent identification of the extracted 18 N-terminal amino acids as being identical to the N-terminal part of the *neuroglian* cDNA clone (Bieber et al., 1989).

The Neurotactin antibody labels secondary neurons at different *D. melanogaster* life stages. It was raised against the first 280 N-terminal amino acids of the *neurotactin* gene (Hortsch et al., 1990). This monoclonal antibody detected the same pattern as two alternative antibodies (Hortsch et al., 1990; Piovant & Lena, 1988). Neither Neuroglian nor Neurotactin antibodies showed signal in *T. castaneum* (data not shown).

### Imaging

All *T. castaneum* imaging data was obtained using a ZEISS LSM 980 (Oberkochen, Germany). Most of the data was acquired using a 40x Gycerol immersion objective (LD LCI Plan-Apochromat 40x/1.2) with 405 nm and 488 nm Diode lasers and 561 nm DPSS laser as well as a GaAsP detection unit and two MA-PMTs. Slice thickness for z projections was always set to optimal. Optical xy resolution was between 1024×1024 to 2048×2048, and always at an 8-bit depth. Scanning was bidirectional, speed between 5-8 and line averaging at 2. *D. melanogaster* data was obtained with a ZEISS LSM 700 Imager M2 (Oberkochen, Germany) using a 40x OIL objective (40x/1.3) and sections were 1.2 to 2 µm thick.

### Data analysis

All imaging analysis was performed in FIJI (Schindelin et al., 2012; http://fiji.sc/). For the 3D models, stacks of digitized images of confocal sections were imported into the 3D modeling TrakEM2 plugin in FIJI software (Cardona et al., 2012). Digital atlas models of three types of structures, including neuropil compartments, neuropil fascicles, and cortical fibre tracts, were created. Compartment boundaries, recognisable by discontinuities in synapse densities and fibre texture, were manually outlined on consecutive sections as “area lists” from which polygonal surface models were calculated. Fascicles and tracts were captured manually as Bezier curves and rendered by the TrakEM2 software as elongated cylinders (“pipes”) of adjustable diameters.

Adobe Photoshop was used to assemble and annotate confocal sections as well as 3D models and Adobe Illustrator was used to draw brain sketches.

## Results

### 1. Overview of the terminology describing insect brain development

Building blocks of the insect brain are (1) the cell bodies (somata) of neurons which form an outer layer (cortex or rind); (2) long axons extended by the cell bodies towards the inner neuropil; (3) terminal branches (axons and dendrites with synapses) within the neuropil; (4) glial cells surrounding somata, as well as the surface of the cortex and neuropil (Figure 1A). Importantly, all cell bodies that stem from one neural stem cell (neuroblast) remain in close proximity within the cortex and the axons of these groups of neurons bundle together to form tracts, or fascicles (Hartenstein et al., 2021). Cells and projections derived from one neuroblast constitute a neural lineage. The basic pattern of fascicles remains constant throughout development, thereby representing an anatomical scaffold that is visible from the larva to the adult. Large parts of the scaffold appear to be largely conserved among different insects, as shown for representative fascicles in Figure 1C-E. Glial processes and fascicles demarcate boundaries between discrete anatomical units (compartments or neuropil domains) of the neuropil (Figure 1A). We will in the following section provide an overview of the main neuropil fascicles and compartments, previously defined and annotated for *D. melanogaster* (Ito et al., 2014; Lovick et al., 2013; Pereanu et al., 2010) and a number of other species (Bressan et al., 2015; Immonen et al., 2017; Figure 1E-F; see Table 2 and 3 for terminology of fascicles), that were used in this work to analyse the structure and development of the *T. castaneum* brain.

**Table 3:**
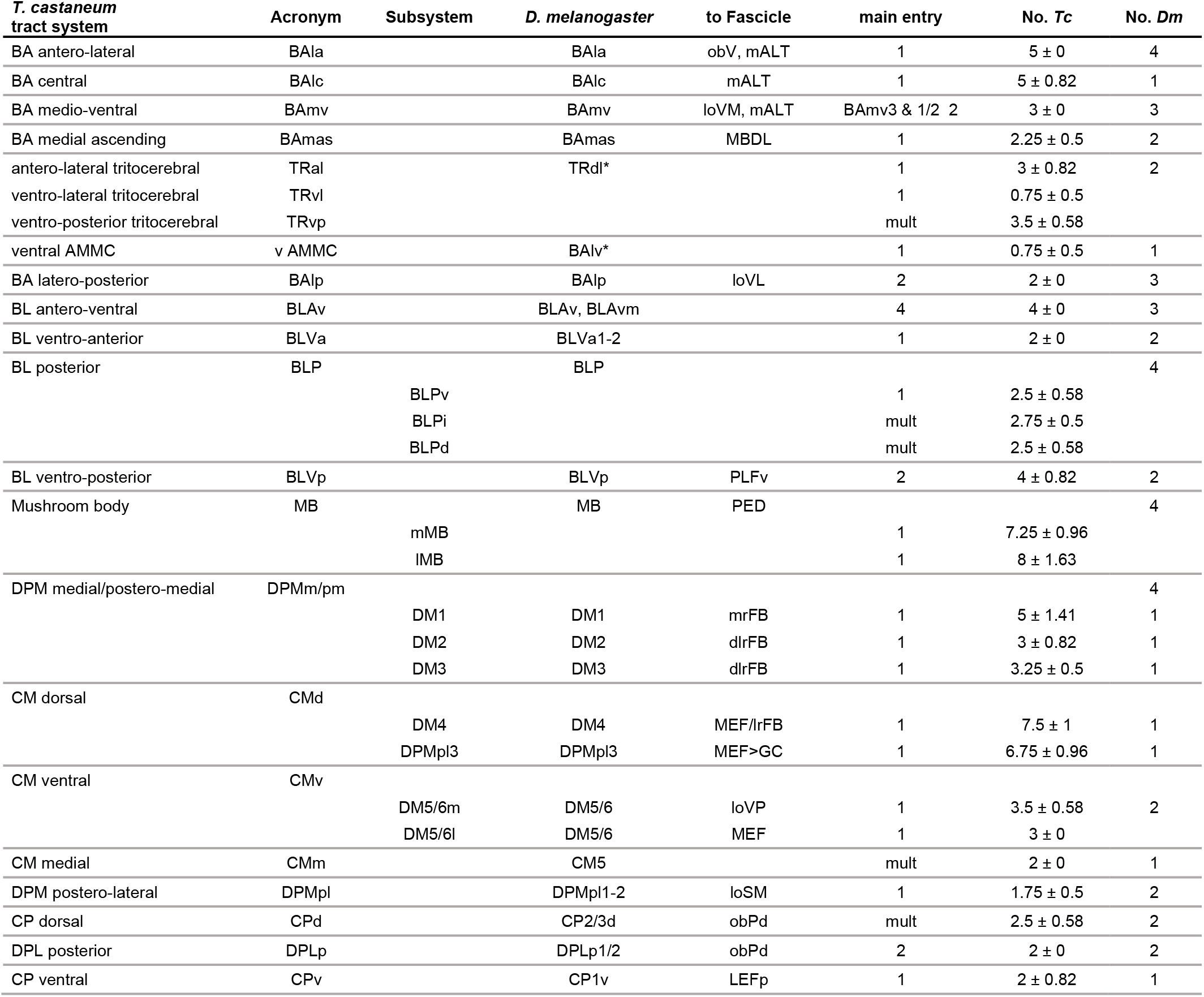
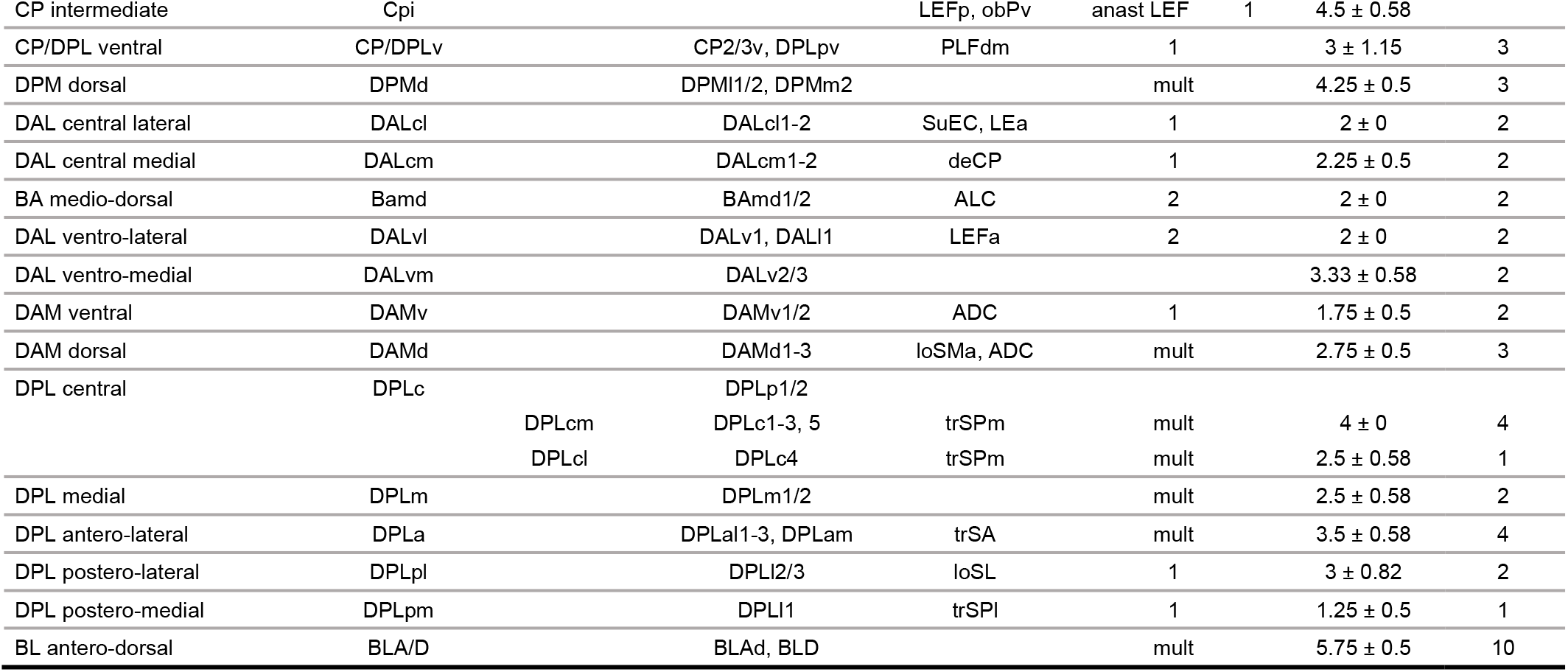
Tract naming in *T. castaneum* and comparison to *D. melanogaster*, with respect to naming, fascicle entry point and number. Numbers of tracts for *T. castaneum* are means with standard deviation of a N=3 population.

It is essential to note that the orientation of the brain within the head capsule differs strongly in insect species (Figure 1B). The *T. castaneum* brain is bent posteriorly, distorting the relationship of body axis and neuraxis in cross-species comparisons. This can make descriptions of anatomy cumbersome when keeping true to the body axis, or confusing when trying to adhere to the neuraxis. For sake of comparability with the *D. melanogaster* atlas, we decided to treat the *T. castaneum* orientations “as if” they were like in the *D. melanogaster* brain (this converts the neuroposterior position from the *T. castaneum* to a *D. melanogaster* ventral position, i.e. “NP=’V’”, Figure 1B).

The insect brain includes two major subdivisions, the supraesophageal ganglion and subesophageal ganglion. Both “ganglia” are complex structures that emerge from the fusion of several “true” (=segmental) ganglia, called neuromeres. The subesophageal ganglion (not further considered in the present work) is composed of three neuromeres (mandible, maxilla, labium); the supraesophageal ganglion is also divided into three units, tritocerebrum, deutocerebrum, and protocerebrum (Figure 1B, C). Neuromeres are most easily defined in the embryo, where discrete clusters of neuroblasts can be assigned to each neuromere (*Tenebrio molitor*: Urbach et al., 2003; *D. melanogaster*: Urbach & Technau, 2003a, 2003b; Younossi-Hartenstein et al., 1996). The axon bundles of neural lineages predominantly innervate the corresponding neuromeres (Kumar et al., 2009). In addition, discrete peripheral nerves are associated with each neuromere. The sensory elements of these nerves form ventrally located centres in the brain. The pharyngeal nerve (innervating the mouth cavity and pharynx) targets the ventral tritocerebrum and the antennal nerve (Kendroud et al., 2018). Based on the upward tilt of the neuraxis, these sensory centres occupy an anterior position. The remainder of the supraesophageal ganglion belongs to the protocerebrum. It consists of the central brain, including mushroom body and central complex, and the optic lobe, which processes input from the compound eye (Figure 1C).

### 2. Compartments and Fascicles of the *Tribolium* larval brain

#### 2.1. Deutocerebrum and tritocerebrum

##### 2.1.1. Antennal lobe (AL; ventral deutocerebrum)

The AL is a spherical compartment that tips the anterior-ventral neuropil surface and has an internal glomerular structure (larva: Figure 2A1, B1, C1; adult: Figure 3A1, B1, C1). It is innervated by the antennal nerve (AN) which enters the AL at its ventro-anterior-lateral edge (larva: Figure 2B1, C1; adult: Figure 3B2). The AL is bounded on all sides, except posteriorly, by a glial layer (Figure 2A1); at its posterior surface the AL borders the ventromedial cerebrum (VMC; dorsal deutocerebrum) and the antenno-mechanosensory motor centre (AMMC; larva: Figure 2A3, C2, D2; adult: Figure 3C3). Left and right antennal lobes are interconnected by the large antennal lobe commissure (ALC; Ito et al., 2014; larva: Figure 2B2, D2, D3; adult: Figure 3B2, C3). The loss of glomerular texture, as well as the fan of antennal projection neurons (see below) that leave the AL towards dorso-posteriorly as the medial antennal lobe tract (mALT; Ito et al., 2014) marks the transition between AL and VMC. The mALT is one of the most voluminous neuropil fascicles described for most insect brains. Leaving the AL it projects posterior dorsally, passing underneath the mushroom body medial lobe (ML), then the central complex (CX), to reach the mushroom body calyx (CA; larva: Figure 2B2-4, D1-3; adult: Figure 3B1-4, C1, C3). Here axons turn laterally to end in the lateral horn (LH; Figure 2D4).

**Figure 2:**
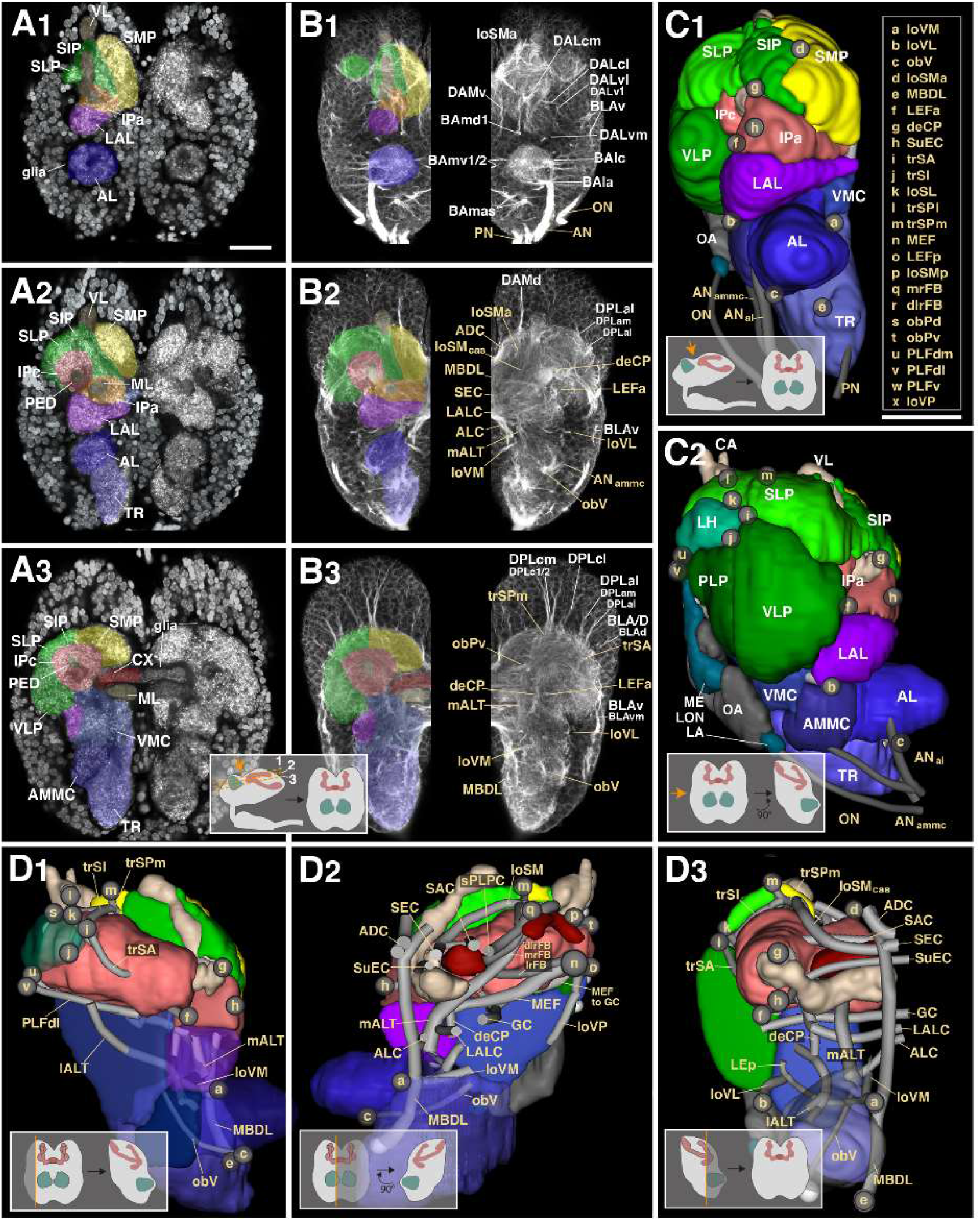
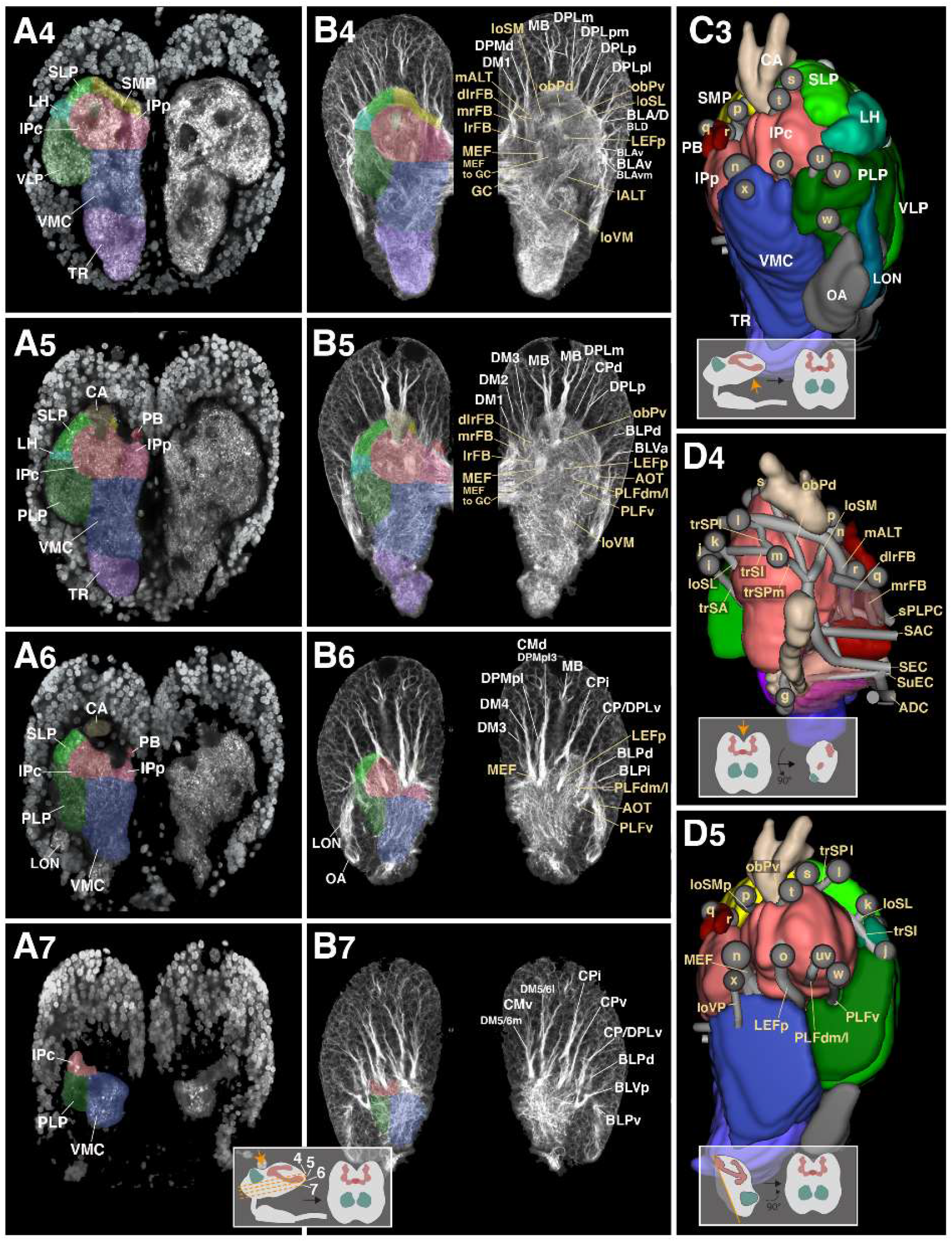
Structure of the early larval brain of *T. castaneum*: Neuropil compartments and fascicles. **(A1-7)**: Z-projections of frontal confocal sections of a larval brain hemisphere labeled with an antibody against synapsin (neuropil in brain center) and with DAPI (neuronal nuclei in surrounding brain cortex). Z-projections represent brain slices of approximately 8–10mm thickness and are arranged in anterior (A1) to posterior (A7) sequence at levels shown in inset at bottom of A7. The left side of each panel has neuropil compartments shaded in different colors and annotated with white letters; right sides show mirrored hemispheres without coloring. (B1-7): Z-projections of frontal sections of larval brain hemisphere labeled with Tubulin antibody, highlighting neuronal cell bodies, tracts and fascicles. Panels are constructed as explained for the A1-7 series and show sections at antero-posterior levels corresponding to those of opposite panels of this series. Compartments are color-coded as in (A1-7) on left side of panels. Right sides show mirrored hemispheres in which fascicles are annotated in beige letter, and axon tract systems in white letters. (C1-3): Digital 3D models of one brain hemisphere in anterior view (C1), lateral view (C2) and posterior view (C3). For orientation see insets at bottom of these panels. Neuropil compartments are rendered in the same colors as in sections shown in (A1-7) and (B1-7) and annotated in white lettering. Small spheres annotated in beige single letters represent locations where neuropil fascicles (themselves not visible in these surface views) begin. For correspondence of single letters with fascicle names see inset at the right of C1. (D1-5): Digital 3D models of one brain hemisphere in lateral view (D1), medial view (D2), anterior view (D3), dorsal view (D4) and posterior view (D5; see insets at bottom of panels for orientation). As in series (C1-3) compartments are color coded, but parts of compartments facing the viewer are cut away to allow observing fascicles (represented as gray tubes) which otherwise would be hidden from view. Starting points of fascicles are shown has single letter-containing spheres; fascicles themselves are annotated in beige lettering. For all abbreviations see Table 2. Scale bars: 25µm (A1-7, B1-7; C1-3, D1-5).

**Figure 3:**
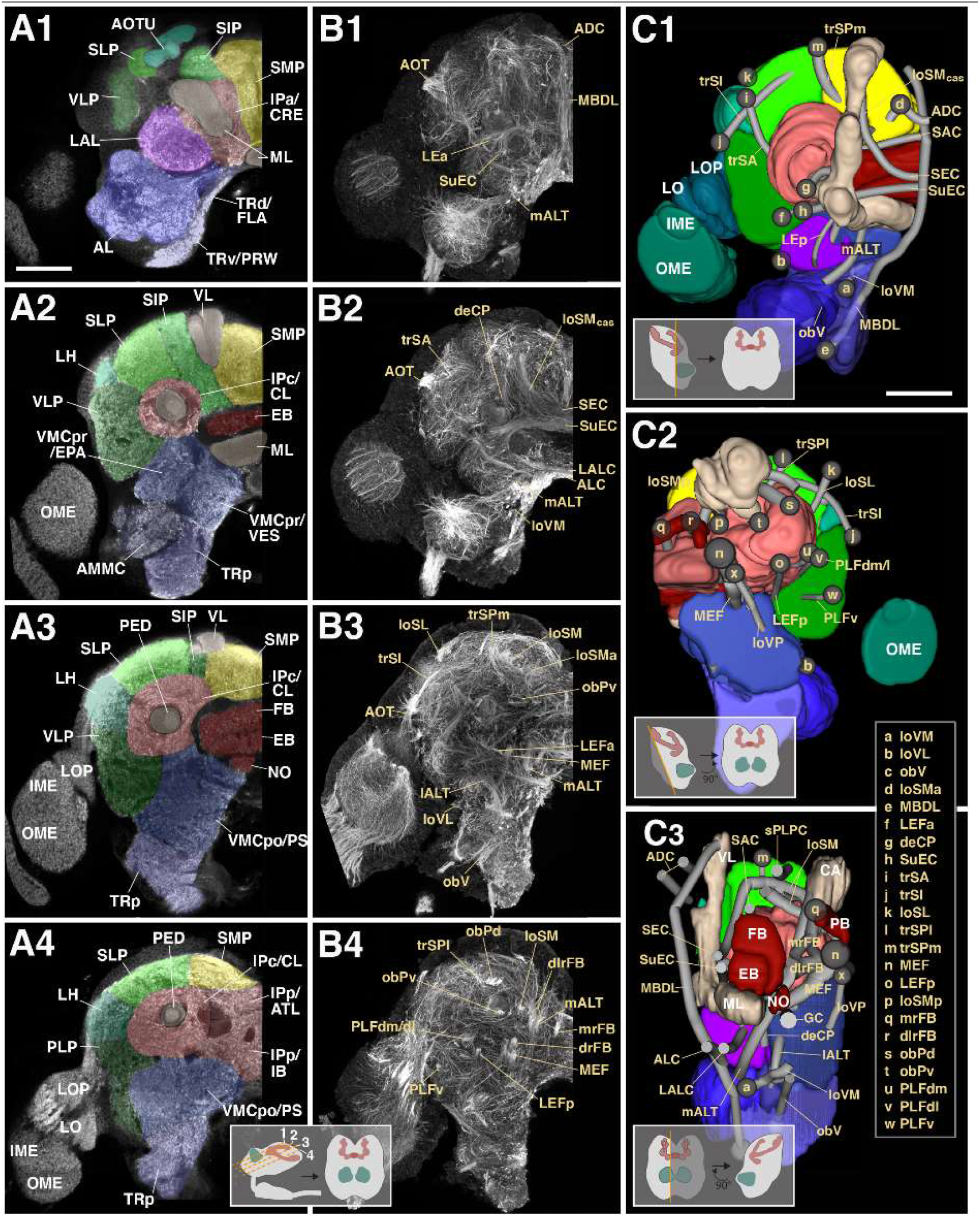
Structure of the adult brain of *T. castaneum*: Neuropil compartments and fascicles. (A1-4): Z-projections of frontal confocal sections of adult brain hemisphere labeled with an antibody against synapsin. The four z-projections represent brain slices of approximately 8–10mm thickness and are arranged in anterior (A1) to posterior (A4) sequence at levels shown in inset at bottom of A4. Compartments are annotated with white letters. (B1-4): Z-projections of frontal sections of larval brain hemisphere labeled with Tubulin antibody, arranged as explained for the A1-4 series. Fascicles are annotated in beige lettering. (C1-3): Digital 3D models of one brain hemisphere in anterior view (C1), posterior view (C2) and lateral view (C3). As explained for Figure 2 D1-5, compartments are color coded, but parts of compartments facing the viewer are cut away to allow for observing fascicles. Starting points of fascicles are shown has single letter-containing spheres; fascicles themselves are annotated in beige lettering. For all abbreviations see Table 2. Scale bars: 50µm (A1-4, B1-4; C1-3).

A second pathway linking the AL as well as the adjacent AMMC to the superior protocerebrum is the lateral antennal lobe tract (lALT; Ito et al., 2014). In *D. melanogaster*, this bundle, formed by subsets of neurons of the lineage BAlc/ALl1 (Ito et al., 2014; Wong et al., 2013; Yu et al., 2013) is very thin and does not appear clearly labelled by global markers like tubulin or BP104. By contrast, the corresponding fibre system forms a thick, conspicuous fascicle in *T. castaneum* from L1 to adult (as also reported for the dung beetle, *S. satyrus*; Immonen et al., 2017). Fibres leaving the posterior surface of the AL, and medial surface of the AMMC, converge into a massive bundle that projects dorso-laterally, passing in between the ventrolateral protocerebrum and inferior protocerebrum (see below) and reaching the lateral horn (larva: Figure 2B4, D1, D3; adult: Figure 3B3).

##### 2.1.2. Anterior periesophageal neuropil (Tritocerebrum)

The tritocerebrum (TR; called periesophageal neuropil, PENP in Ito et al., 2014, and a recent study of the coleopteran *S. satyrus*; Immonen et al., 2017) forms a neuropil domain that is located ventrally and medially of the AL. As in *D. melanogaster* (Hartenstein et al., 2017; Rajashekhar & Singh, 1994) we distinguish a ventral tritocerebrum [TRv; also called Prow (PRW) in Ito et al., 2014] that forms the very anterior tip of the SEZ, and a dorsal tritocerebrum [TRd; also called flange (FLA) in Ito et al., 2014] that borders the wall of the esophageal foramen. We termed the large part of the tritocerebrum that continues all the way to the posterior surface of the brain posterior tritocerebrum (TRp). The entry of the thick pharyngeal nerve (PN) that carries gustatory/mechanosensory afferents from the external and internal mouthparts in *D. melanogaster* and other insects; Kendroud et al., 2018) enters the TRv at its anterior ventral surface (larva: Figure 2B1, C1; adult: Figure 3B2).

Two ventral longitudinal fascicles demarcate the border between the TR and the adjacent deutocerebrum: (1) The longitudinal ventromedial fascicle (loVM; Pereanu et al., 2010; Lovick et al., 2013; IFS in the adult fly and beetle brain; Ito et al., 2014; Immonen et al., 2017) begins in a depression of the anterior neuropil surface in between AL (laterally) and TR (medially; larva: Figure 2B2-5, D1-3; adult: Figure 3B2, C1, C3). The bundle continues posteriorly along the lateral TR border. It splits into several branches that reach into the ventromedial cerebrum (VMC; see below). One highly conspicuous branch, called posterior lateral ellipsoid fascicle (LEp; Lovick et al., 2013) curves dorsally, first following a lateral course, then bending towards medially and dorsally through the lateral accessory lobe (LAL) towards the central complex (larva: Figure 2D3; adult: Figure 3C1). (2) The oblique ventral tract (obV; not previously named): begins near the entry point of the antennal nerve into the AL and extends postero-medially along the ventrolateral border between TR and VMC (larva: Figure 2B2–3, D1-3; adult: Figure 3B3, C1, C3).

##### 2.1.3. Antenno-mechanosensory and motor centre (AMMC)

The AMMC is located posteriorly of the AL and laterally of the TR. It is innervated at its anterolateral surface by a posterior branch of the antennal nerve (ANp; Figure 2B2, C1-2) that carries sensory afferents of mechanoreceptors located on the antenna (Kamikouchi et al., 2006; Sant & Sane, 2018). Aside from the fact that the AMMC does not have a glomerular structure, its borders with the AL (anterior) and VMC (posterior-medially) are difficult to define. In accordance with compartment definitions in *D. melanogaster* we took the obV tract as a marker delineating the border between AMMC (dorso-laterally of obV), VMC (see below; dorso-medially of obV), and tritocerebrum (ventromedially of AMMC; Figure 2B2-B3, D3). At the neuropil surface, a conspicuous indentation demarcates the AMMC from the ventrally adjacent tritocerebrum (larva: Figure 2A3; adult Figure 3A2).

##### 2.1.4. Ventromedial cerebrum (VMC; dorsal deutocerebrum)

The neuropil domain located posteriorly of the antennal lobe, and dorsolaterally of the tritocerebrum constitutes the VMC. Posteriorly the VMC reaches the neuropil surface; this region, in the adult fly and beetle brain, is called the posterior slope (PS; Ito et al., 2014; larva: Figure 2B4-7; C3, D2, 3, 5; adult: Figure 3A2-4, D2-3). Postero-laterally, a deep indentation of the neuropil surface demarcates the VMC from the ventrolateral and posterior lateral protocerebrum (VLP, PLP; Figure 2A4-7, D5; adult: Figure 3A3-4, C2). Additional VMC subdivisions, defined for the *D. melanogaster* and *S. satyrus* adult brain (Ito et al., 2014; Immonen et al., 2017) and shown here for the adult *T. castaneum* brain, include the vest (VES), epaulette (EPA), and gorget (GOR; Figure 3A2-3).

Neuropil fascicles delineating the boundaries of the VMC are:

(1, 2) The loVM and obV as markers of the anterior and medial boundaries between VMC, AL and TR were introduced above. (3) Medial antennal lobe tract (mALT): Dorsoanteriorly, the VMC contacts the lateral accessory lobe (LAL; see below). For the most part, the border between these two compartments is formed by the fan of fibres that converge upon the mALT (larva: Figure 2B2, D2; Figure 2B2, adult: Figure 3C1, 3). (4) descending protocerebral tract (deCP; Pereanu et al., 2010; Lovick et al., 2013): Another easily visible tract that begins at the anterior neuropil surface, laterally adjacent to the base of the VL (larva: Figure 2B2, C1; adult: Figure 3C1). The tract sweeps posterior medially, encircling the VL, and then, after passing the peduncle, turns straight ventrally (larva: Figure 2B3, D3; adult: Figure 3B2, C1). This descending leg of the deCP defines a vertical plane that demarcates the transition of LAL (antero-dorsally of the plane) into VMC (posterior of the plane (Figure 2D2, 3C3). (5) Great commissure (GC): At a posterior level, the great commissure (GC; Ito et al., 2014; Immonen et al., 2017) which interconnects ventral neuropil domains of the left and right brain hemisphere, extends closely underneath the dorsal border of the VMC (larva: Figure 2B4, D2-3; adult: Figure 3C3). (6) Longitudinal ventrolateral fascicle (loVL; Pereanu et al., 2010; Lovick et al., 2013): The loVL forms a bundle that enters from anteriorly into the cleft between VLP, AMMC and VMC (larva: Figure 2B1-3, D3; adult: Figure 3B3, C1).

#### 2.2. Protocerebrum

The protocerebrum accounts for most of the volume of the insect brain. It is customary to distinguish the easily recognizable “structured” neuropil compartments, represented by the central complex (CX), mushroom body (MB), and optic lobe (OL), from the so-called unstructured domains that comprise the ventral (or basal) protocerebrum, inferior protocerebrum, and superior protocerebrum. The inferior protocerebrum (IP) surrounds the mushroom body. The superior protocerebrum and ventral protocerebrum flank the IP dorsally and ventrally, respectively (Ito et al., 2014; Immonen et al., 2017). The superior protocerebrum is further divided into superior medial protocerebrum (SMP), superior lateral protocerebrum (SLP), and superior intermediate protocerebrum (SIP), and lateral horn (LH). The ventral protocerebrum includes the ventrolateral protocerebrum (VLP) and lateral accessory lobe (LAL).

##### 2.2.1. Central complex, mushroom body and optic lobe

The central complex, mushroom body and optic lobe are landmark structures of the insect brain; they are easily recognizable because they are set apart from the surrounding neuropil by a continuous glial sheath (larva: Figure 2A1-3, B1-3; adult: Figure 3A1-4). The central complex consists of the protocerebral bridge (PB), the fan-shaped body (FB; previously: upper division of the central body), ellipsoid body (EB; formerly: lower division of the central body), and noduli (NO; previous description in Dreyer et al., 2010; Farnworth et al., 2020). The latter two are only recognizable in the adult (compare Figure 2A3 and Figure 3A3). The PB forms a thin, arched compartment attached to the posterior surface of the protocerebrum (larva: Figure 2A6, C3; adult: Figure 3C2-3). Large fascicles associated with the central complex are the roots of the fan-shaped body, the lateral ellipsoid tracts, and three commissures. The roots of the fan-shaped body (Farnworth et al., 2020; Hartenstein et al., 2015; Lovick et al., 2013; Pereanu et al., 2010) are formed by the columnar neurons that interconnect thin “slices” (columns) of the compartments that make up the CX, i.e., the PB, FB, and EB. Columnar neurons derive from four large (“type 2”) lineages DM1-DM4, with DM1 located furthest medially/anteriorly, and DM4 most laterally/posteriorly (Farnworth et al., 2020; see below). Somata of the columnar neurons fill the dorsomedial cortex of the protocerebrum (“pars intercerebralis”). Fibres destined for the CX converge upon the cleft between the PB and adjacent IPp. They collect in three bundles, the lateral root (lrFB; tract of DM4; also called “W” tract in the classical literature), dorso-lateral root (dlrFB; DM3 and DM2; tracts “X” and “Y”) and medial root (DM1; tract “Z”; larva: Figure 2B4-5, C3, D2; adult: Figure 3B4, C3). Reaching the FB the roots split up into a plexus of decussating bundles, called the posterior plexus of the fan-shaped body (pplFB; Lovick et al., 2013; Figure 4B4).

**Figure 4:**
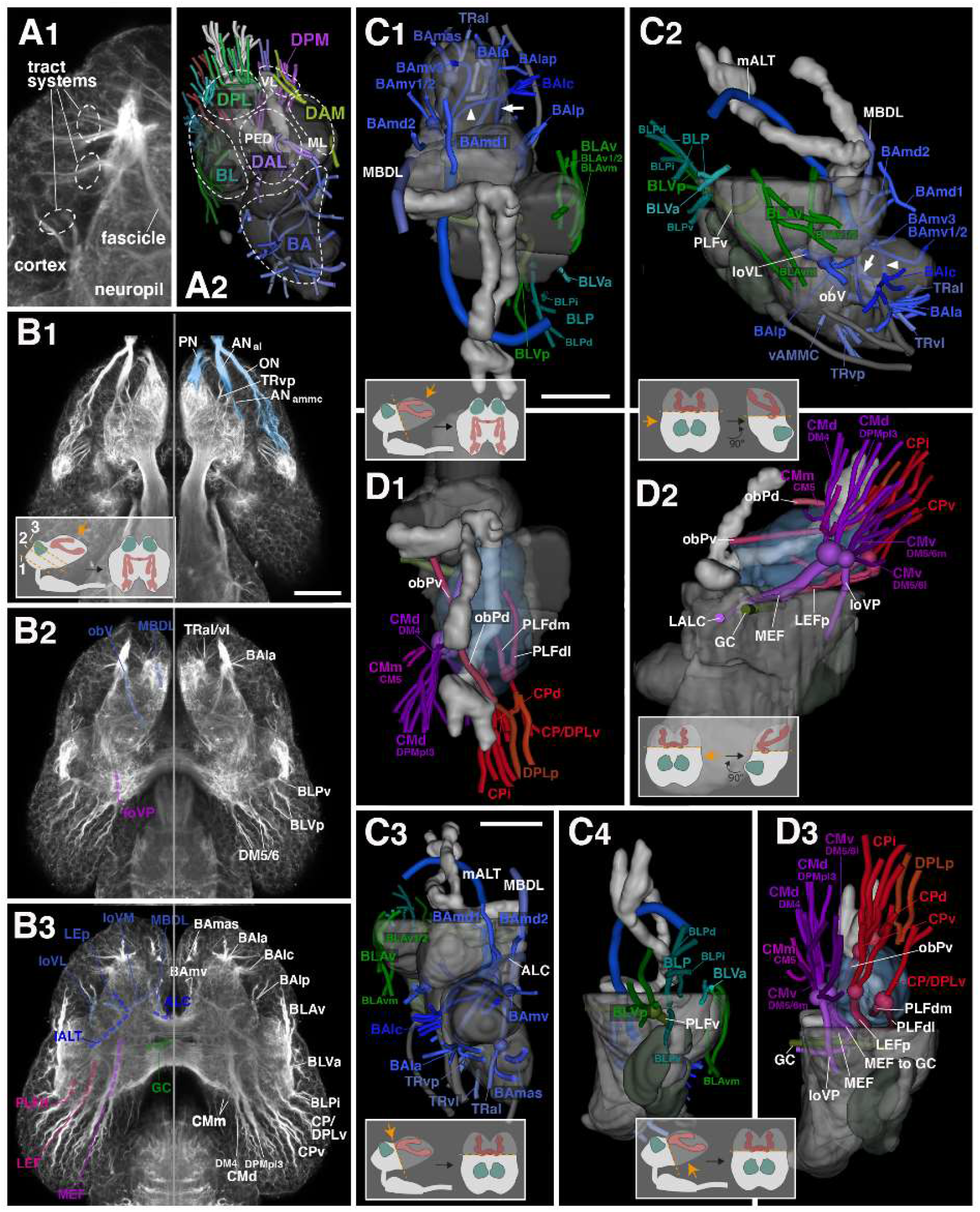
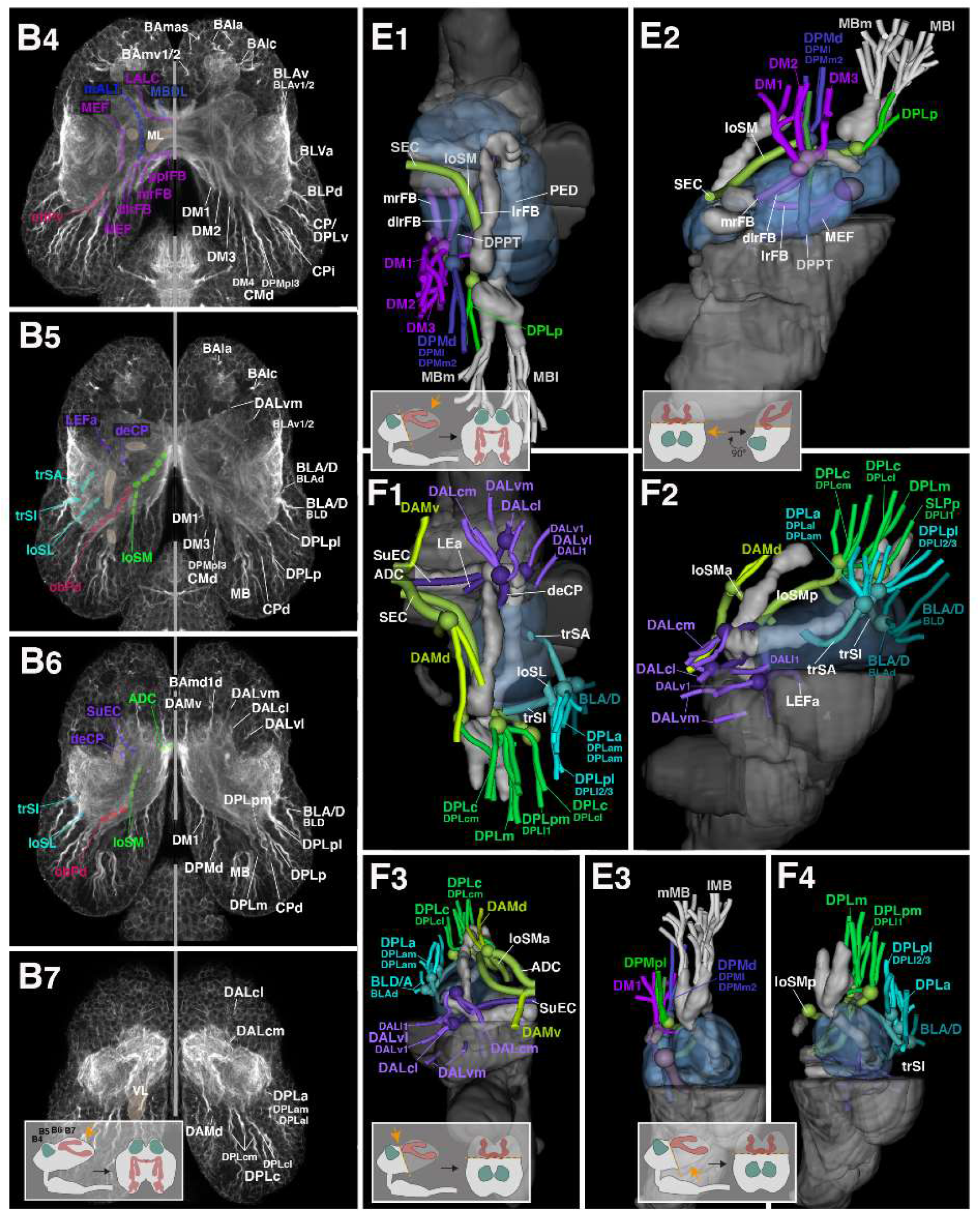
Structure of the early larval brain of *T. castaneum*: Axon tract systems in relation to fascicles. (A1): Confocal section of larval brain tissue labeled with Tubulin antibody, demonstrating appearance of tract systems in brain cortex, and fascicles resulting from confluence of tracts in neuropil. (A2): 3D digital model of larval brain hemisphere in antero-lateral view. Neuropil surface is rendered as semi-transparent gray sheet. Mushroom body (ML medial lobe; PED peduncle; VL vertical lobe) is rendered opaque light gray. Axon tracts entering neuropil are shown in their respective colors used throughout all panels of Figures 4 and 5. Hatched lines encircle super-sets of axon tracts named according to their topology relative to the mushroom body (for detail, see text). (B1-7): Z-projections of horizontal sections of larval brain hemisphere labeled with Tubulin antibody. Panels show sections at ventral (B1) to dorsal (B7) levels as indicated in insets at the bottom of (B1) and (B7). Right and left sides show mirrored hemispheres; on left sides, fascicles are shaded by hatched lines, and annotated, in colors used throughout all panels of Figure 4. White lettering on right sides annotates tract systems. Right side of (B1) shows peripheral nerves entering brain in blue shading. (C1-4, D1-3, E1-3, F1-4): Digital 3D models of one hemisphere in dorsal view (C1, D1, E1, F1), lateral view (C2, F2), medial view (D2, E2), anterior view (C3, F3), and posterior view (C4, D3, E3, F4). Neuropil surface is rendered as semi-transparent gray sheet, mushroom body is shown in light gray. Neuropil dorsal of the peduncle is cut away, except for the central inferior protocerebrum, shown in cyan in panels (D1-3, E1-3, F1-4). Reconstructions of axon tracts appear as tubes entering the neuropil in close association to fascicles. The coloring scheme follows the one used for the fascicles. The four sets of panels (C, D, E, F) show subsets of axon tract systems ordered by topology. (C1-4) Systems associated with the deuterocerebrum and tritocerebrum (BA, TR: blue) and ventrolateral protocerebrum (BL: green). (D1-3) Systems entering the posterior surface of the protocerebrum (CM: purple; CP: red). (E1-3) Systems of the dorso-medial protocerebrum (DM/DPM: magenta/dark blue; DPMpl: green; MB: white). (F1-4) Systems of the dorso-lateral protocerebrum (DAL: lilac; DPL: green/cyan) and anterior dorso-medial protocerebrum (DAM: light green). Tract systems are annotated in their respective colors; fascicles are annotated in white. For all abbreviations see Table 2. Scale bars: 25µm (B1-7; C1-2, D1-2, E1-2, F1-2; C3-4, D3, E3, F3-4).

Fibre bundles entering the central complex from laterally include the anterior and posterior lateral ellipsoid fascicles (LEa, LEp; Lovick et al., 2013). The LEp, a branch of the loVM, was described above (see section 2.1.2). The LEa is formed by fibres that enter from the antero-lateral cortex, pass underneath the ML at its junction with the VL, and continue posteromedially towards the central complex (larva: Figure 2D2, D3; adult: Figure 3B1, B2). Closely adjacent to the entry of the LEa are fibres that project medially across the midline, forming the subellipsoid commissure (SuEC; Pereanu et al., 2010; Lovick et al., 2013; larva: Figure 2B2, D3; adult: Figure 3B1, B2, C1, C3). This conspicuous commissure lies in between ML (anteriorly) and central complex (posteriorly). Crossing the central complex slightly more dorsally is the supraellipsoid commissure (SEC; Ito et al., 2014; larva: Figure 2B2, D3; adult: Figure 3B1, B2, C1, C3); further posteriorly, covering the “roof” of the central complex, is the superior arch commissure (SAC; Ito et al., 2014; larva: Figure 2D2, D3; adult: Figure 3C1, C3).

The MB consists of a large number of neurons (Kenyon cells) with cell bodies in the posterior cortex (previous description for adult: Dreyer et al., 2010). Dendritic branches of the Kenyon cells form the calyx at the dorsoposterior neuropil surface (larva: Figure 2A5-6, C3, D4-5; adult: Figure 3C2). Bundled Kenyon cell axons continue straight anteriorly as the peduncle (PED), which projects forward all the way to the antero-lateral brain surface (larva: Figure 2D2; adult: Figure 3C3). Here the PED bifurcates into the medial lobe (ML) and vertical lobe (VL). The VL is directed straight dorsoposteriorly, extending into the cortex (larva: Figure 2A1-2, D2; adult: Figure 3A2-3, C3). The ML projects posteromedially, reaching the midline right in front of the central complex (larva: Figure 2A3, D2; adult: Figure 3A2, C1, C3).

The optic lobe (OL), which receives input from the stemmata of the larva and later the compound eye of the adult, protrudes from the posterolateral surface of the protocerebrum. In the adult, the compartments of the OL (from distal to proximal: lamina, medulla, lobula, lobula complex) are easily delineated (Dreyer et al., 2010; Figure 3A2-4, C1). During larval stages, rudiments of these compartments exist, embedded into strands of neuroepithelial progenitor cells (optic anlagen; OA; Figure 2B6, C3). Lamina and medulla are set apart; medulla and lobula complex of the early larva appear as a single conical process (larval optic neuropil, LON) attached to the posterior lateral protocerebrum (Figure 2A6, B6, C3).

##### 2.2.2. Inferior protocerebrum (IP)

We distinguish a central (IPc), anterior (IPa) and posterior domain (IPp) of the IP. All of these are closely associated with the peduncle and lobes of the mushroom body.

*Central domain of the inferior protocerebrum (IPc)*: The IPc forms a hollow cylindrical structure that surrounds the MB peduncle in the centre of the brain hemisphere. Due to this configuration the IPc of the adult fly brain was dubbed “clamp” (Ito et al., 2014; for dung beetle: Immonen et al., 2017). Borders to the adjacent superior protocerebrum, ventral protocerebrum, and posterior inferior protocerebrum are defined by several longitudinally and transversally oriented fascicles. Among the former we distinguish the medial equatorial fascicle (MEF), longitudinal ventroposterior fascicle (loVP), longitudinal superior medial fascicle (loSM), lateral equatorial fascicle (LEF), posterior lateral fascicle (PLF), and oblique posterior fascicle (obP).

(1) The MEF (Ito et al., 2014) is a thick bundle that originates at the posterior neuropil surface, laterally adjacent to the protocerebral bridge (larva: Figure 2B4-6, C3, D2, D5; adult: Figure 3A3-4, C2-3). It is formed mainly by anteriorly directed axons of multiple lineages located in the posterior cortex (see below). The MEF gives off a branch into the great commissure (larva: Figure 2B4-5), and then extends forward towards the LAL. Here it turns medially to cross into the contralateral LAL, forming part of the LAL commissure (LALC; Ito et al., 2014; larva: Figure 2D2-3; adult: Figure 3B2, C3).

(2) The loVP (Pereanu et al., 2010; Lovick et al., 2013) originates at a position directly ventral of the MEF (larva: Figure 2C3; adult: Figure 3C2). It turns ventrally, extending near the posterior surface of the VMC compartment (larva: Figure 2D2, D5; adult: Figure 3C2-3).

(3) The loSM (Pereanu et al., 2010; Lovick et al., 2013; also called SFS in adult; Ito et al., 2014; Immonen et al., 2017) starts at the posterior neuropil surface dorsally and laterally of the MEF (larva: Figure 2C3; adult: Figure 3C2). It consists of multiple bundles directed upward and forward along the dorsomedial border of the IPc compartment (larva: Figure 2B4, D2-4; adult: Figure 3B2-4, C2-3). Reaching the level of the VL most components of the loSM bend ventromedially (“cascade” of the loSM) and, in part, cross the midline at a level dorsally adjacent to the central body, forming the supraellipsoid commissure (see above).

(4) The posterior LEF (LEF; Ito et al., 2014) also begins at the posterior neuropil surface, ventrally of the calyx, and follows a straight anteriorly directed course parallel to the MEF and peduncle (larva: Figure 2B4-6, C3, D5; adult: Figure 3B4, C2).

(5) The PLF system (Ito et al., 2014) starts out in a depression of the posterior-lateral neuropil surface (larva: Figure 2C3; adult: Figure 3C2). It includes three bundles, called the dorso-lateral, dorso-medial component, and ventral component (PLFdl, PLFdm, PLFv) of the PLF (Lovick et al., 2013). Fibres extend anteriorly along the ventrolateral boundary of the IPc (larva: Figure 2B5-6, D1, D5; adult: Figure 3B4, C2). At an anterior level, the PLFdm turns dorsally, converging on the peduncle; the PLFdl and PLFv remain at a more lateral and ventral level, respectively.

(6) The obP (Pereanu et al., 2010; Lovick et al., 2013) also originates at the posterolateral neuropil surface, close to the calyx (larva: Figure 2C3; adult: Figure 3C2). It projects dorso-medially, passing over the dorsal surface of the peduncle before turning anteriorly, and joining the loSM fascicle (larva: Figure 2B4, D4; adult: Figure 3B4, C2).

In addition to these longitudinal fibre systems, a series of transversally directed bundles that enter the neuropil from dorsally and laterally demarcate the boundary between IPc and superior protocerebrum. They constitute the transverse superior anterior (trSA), transverse superior intermediate (trSI) and transverse superior posterior (trSP) fascicles.

(7) The trSA (Peranu et al., 2010; Lovick et al., 2013; also termed anterior superior lateral protocerebral fascicle, aSLPF, in adult *D. melanogaster* and dung beetle; Ito et al., 2014; Immonen et al., 2017) is a conspicuous crescent-shaped bundle that enters the dorsoanterior protocerebrum, continues on a ventromedial course, followed by a dorsal bend towards the dorsal aspect of the peduncle (larva: Figure 2B3, C2, D3; adult: Figure 3B2, C1). The trSA demarcates the boundary between IPc (ventroposteriorly), superior lateral protocerebrum (SLP; dorsally), and ventrolateral protocerebrum (VLP; ventrolaterally).

(8) The trSI (Peranu et al., 2010; Lovick et al., 2013) is a system of fibres (termed pyriform fascicle, PYF, in the adult fly and dung beetle brain; Ito et al., 2014; Immonen et al., 2017) that originates at the dorso-lateral neuropil surface, posteriorly of the trSA (larva: Figure 2C2, D3-4; adult: Figure 3B3, C2). Fibres project dorsomedially, filling the lower stratum of the superior protocerebrum (dorsally), and demarcating it from the IPc (ventrally).

(9) The trSP (Peranu et al., 2010; Lovick et al., 2013) is formed by fibres that enter the superior protocerebrum dorsally (Figure 2C2, D4; adult: Figure 3B4, C2). One can distinguish a lateral component (trSPl) and a medial component (trSPm). Following entry fibres form loose, crescent-shaped bundles in the lower stratum of the superior protocerebrum directed medially.

*Anterior domain of the inferior protocerebrum (IPa):* The IPa, called crepine (CRE) in the adult brain (Ito et al., 2014; Immonen et al., 2017) forms in essence a hollow cylinder that enwraps the medial lobe (ML) of the mushroom body. However, like the ML itself, the IPa has an uneven, complicated shape. At an anterior level, it wraps around the shaft and proximal part of the ML, leaving its tip (which points posterior-medially) uncovered (larva: Figure 2A1-2, C1-2; adult: Figure 3A1). At the back side of the ML the IPa narrows to cover only the short shaft of the ML (larva: Figure 2A2). Going further posteriorly, the IPa is confluent with the IPc (larva: Figure 2A2). Anterior-ventrally the IPa borders the LAL, and dorsally the SMP (larva: Figure 2C1; adult: Figure 7). The demarcation of IPa from SMP is based on neuropil texture (e.g., density, length, and orientation of tubulin-positive fibres). In particular, one can discern for the SMP, but not the IPa, long fibre arrays swerving around the VL shaft and then turning ventromedially, as well as similarly oriented fibres of the anterior loSM cascade. Separating the IPa from the LAL at the anterior neuropil surface is a distinct groove (larva: Figure C1-2; adult: Figure 7). Along the midline, the massive, vertically oriented median bundles (MBDL; Ito et al., 2014), bidirectional fibre systems connecting SMP and tritocerebrum separate the left and right IPa (larva: Figure 2B2).

*Posterior domain of the inferior protocerebrum* (IPp): this domain, called antler (ANT) and inferior bridge (IB) in the adult fly brain and dung beetle (Ito et al., 2014; Immonen et al., 2017), lies posteriormedially adjacent to the IPc, from which it is demarcated by the MEF and loSM fascicles (larva: Figure 2A4–5, B4-5; adult: Figure 3A4, B4, C2). The protocerebral bridge emerges from the posterior surface of the IPp (larva: Figure 2C3; adult: Figure 3C2). Its ventral border with the VMC is ill defined. Tract systems that enter the IPp and form conspicuous landmarks are the roots of the fan-shaped body (see above).

##### 2.2.3. Superior protocerebrum

The superior protocerebrum forms the dorsal “cap” of the insect brain. Boundaries between the individual neuropil compartments of the superior protocerebrum (SMP, SIP, SLP, LH) appear as furrows at the neuropil surface, distinctive in the in the adult brain, less so at larval stages. Within the neuropil, delineating these boundaries is difficult, because nerve fibres freely cross in between different superior protocerebral compartments at most levels, and synapse density and patterning is similar throughout the superior protocerebrum. Again, virtual planes, defined by distinctive neuropil fascicles and entering tract systems, set the boundaries, as discussed in the following.

*Superior medial protocerebrum:* At an anterior level, the VL of the mushroom body and the anterior cascade of the loSM define the lateral boundary of the SMP (larva: Figure 2A2, B2; adult: Figure 3A2, B2). At an intermediate level, right posteriorly of the VL, the large system of fibres that converges from dorsally onto the neuropil surface and continues as the trSPm (see above) is a defining landmark between SMP (medially) and SLP (laterally; larva: Figure 2B3, D3; adult: Figure 3B3, C1). At its posterior surface, the SMP borders the calyx and the inferior protocerebrum. Several tract systems entering the brain from posteriorly (loSM, roots of the fan-shaped body, see above) delineate the SMP (dorsally) from IPp (ventrally) (IPp; larva: Figure 2A4, B4, D5; adult: Figure 3A4, B4, C2).

*Superior lateral protocerebrum (SLP):* The SLP covers the domain located laterally of the mushroom body VL (anteriorly; larva: Figure A1-2; adult: Figure 7), entry of the trSPm (centrally; larva: Figure 2B3, D3; adult: Figure 3B3, C1) and calyx (posteriorly; larva: Figure 2A6, C3; adult: Figure 7). Towards ventrally, the transverse fascicles (trSA, trSI, trSP; see above) demarcate its borders to the IPa and VLP. The loSL, a bundle formed by fibres entering from the dorsolateral brain cortex (Pereanu et al., 2010; Lovick et al., 2013) defines the border between SLP (anteromedially of loSL) and lateral horn (posteriorly laterally of loSL; larva: Figure 2B4, D5; adult: Figure 3A3, B3, C2).

*Lateral horn (LH):* The LH occupies a posterolateral position within the superior protocerebrum. It receives innervation from terminal fibres of the antennal lobe tracts (mALT, lALT; see above), as well as afferents that ascend from the optic lobe. A landmark tract that develops with the massive postembryonic growth of the optic lobe (and eye) is the anterior optic tract (AOT; Ito et al., 2014); it can be discerned as it exits the lobula complex and turns anteriorly, forming a deep horizontal indentation of the neuropil surface along the boundary between the LH (dorsally) and PLP (ventrally), and, further anteriorly, between VLP and SLP (larva: Figure 2B5-6; adult: Figure 3B2-3). The AOT then turns medially and reaches the anterior optic tubercle (AOTU) that tips the anterior surface of the adult SLP (adult: Figure 3B1). Borders of the LH and adjacent SLP are defined by the loSI (see above).

*Superior intermediate protocerebrum (SIP):* The small domain of the superior protocerebrum that enfolds the mushroom body vertical lobe anteriorly, laterally, and posteriorly is termed SIP (Ito et al., 2014; Immonen et al., 2017). An indentation in the antero-dorsal neuropil surface defines the border between the SIP and laterally adjacent SLP and (in the adult) AOTU (larva: Figure 2A1, B1, C1-2; adult: Figure 3A1). Medially, the fibres of the loSM cascade, as well as the anterior section of the loSM fascicle delineate the SIP from the SMP (larva: Figure 2A2, B2, C1, D3; adult: Figure 3A2, B2). The vertically entering fibres forming the trSPm signify the posterior boundary of the SIP (Figure 2B3, C2, D2; adult: Figure 3B3, C3). Also, the pattern of terminal fibres visible in tubulin-labelled preparations enables to perceive, in particular in the adult, a discontinuity between the neuropil immediately surrounding the VL, and the adjacent SMP and SLP, respectively.

##### 2.2.3. Ventral protocerebrum

*Lateral accessory lobe (LAL):* The LAL is closely connected to the central complex, and a source of output to the motor centres of the ventral nerve cord. The LAL is similar in size and shape to the antennal lobe. Anteriorly and anterior-laterally, the LAL contacts the neuropil surface, forming a conspicuous bulge dorsally of the AL (larva: Figure 2A1-2, B1-2, C1-2; adult: Figure 3A1, Figure 7). The lateral accessory lobe commissure (LALC; Ito et al., 2014) passes in between left and right LAL, at a level dorsoposteriorly of the antennal lobe commissure (larva: Figure 2B2, D2, D3; adult: Figure 3B2, C3). The posterior boundary with the VMC, demarcated by several vertical fibre bundles (e.g., deCP), was discussed above. Posterior-laterally the LAL is separated from the ventrolateral protocerebrum (VLP) by a deep cleft (larva: Figure 2A3, B3, C1-2; adult: Figure 3A1, C1, Figure 7). Dorsally the LAL abuts the inferior protocerebrum (IPa; see above).

*Ventrolateral protocerebrum (VLP) and posterior lateral protocerebrum (PLP):* These compartments form the ventrolateral “bulge” of the brain neuropil. In the adult fly brain, the VLP is further divided into an anterior (AVLP) and posterior (PVLP) subcompartment; the latter is defined by the presence of distinctive, synapse-rich optic glomeruli, formed by the endings of visual afferents from the optic lobe (Ito et al., 2014). AVLP, as well as the PLP bordering the PVLP posteriorly, lack these glomeruli. The PLP is further distinguished as the site of entry of the large number of optic lobe-derived fibre bundles into the central brain. In the fly larva, lacking a differentiated optic lobe, a distinction between PLP, PVLP and AVLP is difficult to make, and the entire complex domain was termed VLP (Pereanu et al., 2010). In contrast, *T. castaneum* exhibits a small, yet differentiated optic lobe with distinct medulla and lobula complex already in the early larva. The lobula complex and bundles of visual afferents originating in the medulla and the complex itself are broadly connected to the posterior protocerebrum, defining this region, which reaches as far forward as the great commissure, as the PLP (larva: Figure 2A5-6, B5-6, C2-3; adult: Figure 3A4, Figure 7). The domain anteriorly adjacent to the PLP (i.e., anterior to the level of the GC) represents the VLP. Optic glomeruli are not evident, so that a clear distinction between PVLPp and AVLP is not possible.

Dorsally, the PLP and VLP border the superior protocerebrum. The points of origin of the trSA and trSI, as well as the anterior optic tract (AOT), demarcate the boundary between PLP/VLP (below) and superior protocerebrum (above; larva: Figure 2A3-5, B3-5, C2, D3-5; adult: Figure 3A2-3, B2-3, C1-2). Anteromedially and posteriorly, deep clefts in the neuropil surface, as well as discontinuous patterning of tubulin positive fibres within the neuropil, delineate the VLP from the LAL, and the PLP from the VMC, respectively (see above). The longitudinal fascicles described in the previous section, PLFm/dl and PLFv, as well as the lateral antennal lobe tract (lALT) demarcate an otherwise ill-defined border between PLP/VLP and medially adjacent IPc (larva: Figure 2A5, B5, D5; adult: Figure 3A4, B4, C2).

### 3. Systems of axon tracts of the larval brain in relationship to compartment and neuropil fascicles

Tubulin labelling acts as global neuronal marker that outlines the cell bodies of the cortex (or cell body rind) and neuronal processes leaving the cell bodies, called axons in the following. Axons converge forming discrete bundles, or axon tracts, that enter the neuropil in a stereotyped pattern. Generally, a pair or small group of tracts enter in close proximity to each other, forming a tract group (tract system; Figure 4A1). Individual members of a tract system could belong to neuron clusters representing different lineages, or parts of lineages (hemilineages, sublineages), which has been definitively shown for the *D. melanogaster* embryo and larva (Larsen et al., 2009; Pereanu & Hartenstein, 2006). Tubulin labelling of axon tracts/tract systems continues within the neuropil, and, in most cases, tracts enter the neuropil fascicles outlined in the previous section. In a number of cases, tracts end shortly after entry and are lost in the diffuse meshwork of tubulin-positive terminal fibres that fill the neuropil. The pattern of tracts can be readily reconstructed in the early larval brain, where (due to the large diameter of the brain cortex) tracts are long and clearly separated from each other. This changes with increasing age of the animal (see following section). We will in the following present the pattern of tract systems observed in the early larva, relating them to the anatomical scaffold provided by the compartments and major fascicles described in the previous section. The data are illustrated in a series of representative z-projections of confocal sections oriented approximately in a horizontal plane (relative to body axis; Figure 4B1-7). In this orientation, most fascicles or tracts are sectioned lengthwise. Confocal images are supplemented with digital 3D models (Figure 4C-F) showing selected subsets of tracts and fascicles. Figure 5 presents 3D models of axon tracts in relationship to neuropil compartments, comparing the pattern observed in *T. castaneum* and *D. melanogaster*.

**Figure 5:**
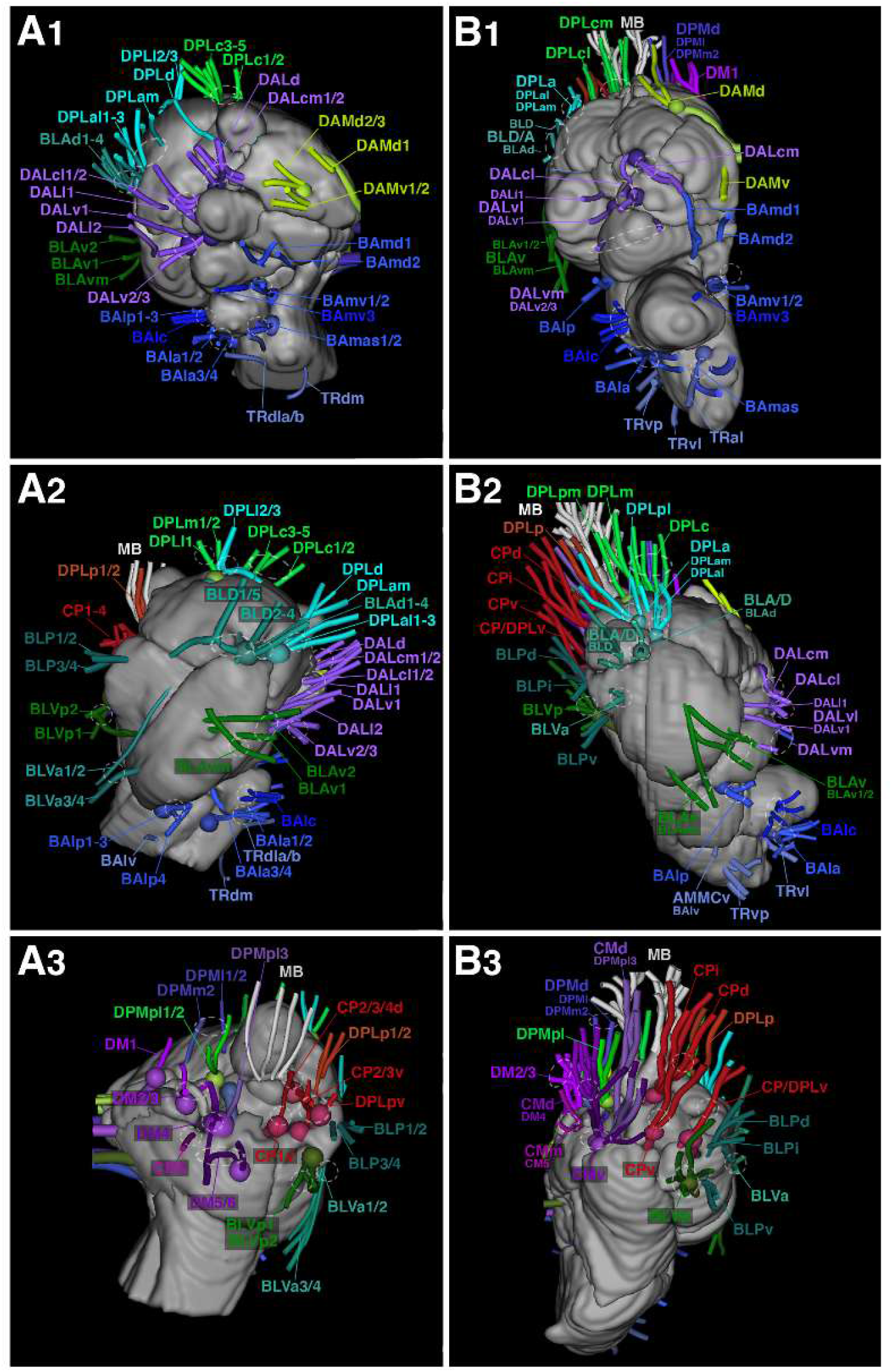
Tract systems of *T. castaneum* in comparison to lineage-associated tracts in *D. melanogaster*. (A1-3, B1-3): 3D digital models of larval (3^rd^ instar) brain hemisphere of *D. melanogaster* (A1-3) and early larval brain hemisphere of *T. castaneum* (B1-3) in anterior view (A1, B1), lateral view (A2, B2) and posterior view (A3, B3). Neuropil surface is shown in gray; axon tracts are rendered and annotated in the same colors as in Figure 4. For all abbreviations see Table 2. Scale bars: 25µm (A1-3; B1-3). *The D. melanogaster* larval model is based on Cardona et al., 2010.

In regard to naming the axon tracts we will adopt the system developed for the *D. melanogaster* larval brain (Lovick et al., 2013; Pereanu and Hartenstein, 2006) in which the neuropil entry point of a tract in relationship to the mushroom body was used as a criterion to define names for super-sets of contiguous tracts, and then further subdivide super-sets into smaller groups, again using position within the super-set as defining feature. In this system, the horizontal plane defined by the peduncle and medial lobes represents the “equatorial” plane that separates dorsal tracts, above the plane, from basal tracts, below. A second, vertical plane defined by the peduncle and mushroom body separates medial from lateral; the plane formed by vertical and medial lobes separates anterior from posterior. Based on this scheme, the super-sets basoanterior (BA), basolateral (BL), dorsoanterior lateral (DAL), dorsoanterior medial (DAM), dorsoposterior lateral (DPL), and dorsoposterior medial (DPM) were established. Two additional super-sets of tracts enter the posterior surface of the neuropil, centered by the entry of the mushroom body peduncle; they were termed centroposterior (CP; entering laterally and ventrally of peduncle) and centromedial (CM; entering medially of peduncle). Tracts entering ventral (i.e., posterior, relative to neuraxis) of the AL were termed tritocerebral (TR).

This overall scheme lends itself acceptably for naming the tracts of the early *T. castaneum* larva (Figure 4A2) and will be followed as a guiding principle. We will group tracts into “systems”, comprised of closely spaced tracts that converge upon a specific neuropil entry position. In cases where tracts could be clearly assigned to the canonical fascicles defined in the previous section, we employed the topology-based nomenclature introduced and consistently used for the *D. melanogaster* larva (e.g., BAla, BAmv). In those cases where an assignment to fascicles is currently not possible, we chose acronyms that respect topology (e.g., DAL, DPL), but that are not part of the series used in *D. melanogaster* (e.g., DPLa for the group including lineages DPLal1-3 and DPLam in *D. melanogaster*).

#### 3.1. Tritocerebrum and Deutocerebrum

##### 3.1.1. Antennal lobe (AL; ventral deutocerebrum)

Aside from the antennal nerve, three tract systems contact the AL:

(1) Entering directly adjacent to the antennal nerve is the basoanterior lateral system (BAla). It encompasses 2 bundles projecting from straight anterior into the AL, and 2-3 bundles located more ventrally and laterally (Figure 2B1; Figure 4B2-3, C1-3). In the neuropil, part of this system continues posteriorly, alongside the AN, and radiates into the root of the antennal lobe tract at the AL/VMC border (Figure 4B3). Other elements of the BAla system form a postero-medially directed bundle that passes ventrally of the AL and continues as the obV fascicle (Figure 4B2, C1-3). In *D. melanogaster*, tracts associated with lineages BAla1-4 closely resemble the BAla system. *D. melanogaster* BAla1 and 2 project into the antennal lobe; BAla1 extends along the antennal lobe tract; BAla3 and 4 target the obV (Figure 5A1-2, B1-2).

(2) The basoanterior central system (BAlc) reaches into the AL from a position lateral and posterior to BAla (Figure 2B1; Figure 4B3, C1-3). 2-3 tracts enter slightly more dorsally and continue straight medially through the AL (Figure 4B3, C1-2: arrowhead), then turn posteriorly to form the major contingent of fibres continuing as the medial antennal lobe tract (mALT), thereby resembling the dorsal hemilineage of *D. melanogaster* BAlc (Figure 5A1, B1). Two additional bundles of the BAlc system contact the AL slightly more ventrally. After entering the neuropil they turn posteriorly, leaving the AL and radiating into the posteriorly adjacent AMMC (Figure 4C1-2, arrow). In the *D. melanogaster* adult, the ventral hemilineage of BAlc innervates the AMMC. We speculate that the ventral contingent of BAlc fibres in *T. castaneum* may correspond to the ventral BAlc hemilineage in flies.

(3) Basoanterior medial ventral system (BAmv): One tract of this system enters the AL from dorsoanteriorly, continues straight through the AL and joins BAla and BAlc elements towards the mALT (Figure 4B5, C1-3). Medially of this tract, which we interpret to represent *D. melanogaster* lineage BAmv3 because it displays the same characteristics (Figure 5A1, B1), are two further tracts that bypass the AL medially and converge into a conspicuous bundle that continues as the loVM fascicle (Figure 2B1; Figure 4B3, C1-3). This pair closely resembles the lineages BAmv1 and 2 in *D. melanogaster* (Figure 5A1, B1).

##### 3.1.2. Tritocerebrum

Four systems of axon tracts are associated with the tritocerebrum. The first two enter at the anterior apex of this compartment, dorsally of the pharyngeal nerve; the other two are located further posteriorly.

(1) Basoanterior medial ascending system (BAmas): A pair of tracts converges from dorsally onto the TR, sharply turns posteriorly and dorsally, and joins the median bundle (MBDL) that flanks the oesophageal foramen and projects to the superior medial protocerebrum (SMP; see previous section; Figure 2B1; Figure 4B3, C1-3). Another component of these tracts, upon entry, projects posterolaterally into the TR. A pair of lineages with equivalent entry and projection are BAmas1/BAmas2 in *D. melanogaster* (Figure 5A1, B1).

(2) Antero-lateral tritocerebral system (TRal): A pair of thinner axon tracts enters laterally of BAmas and remains within the tritocerebral compartment, without connection to the MBDL (Figure 4B2, C3). These tracts may correspond to the TRdl lineage tracts in *D. melanogaster* (Figure 5A1, B1).

(3) Ventrolateral TR system (TRvl): a single tract enters at a level ventrally and laterally of the pharyngeal nerve (Figure 4C2)

(4) Ventroposterior TR system (TRvp): 4-5 tracts enter the TR at its posterolateral surface (Figure 4B1, C2-3). They resemble *D. melanogaster* tritocerebral lineages TRdm, BApd, and BApv, but assigning individual tracts in *T. castaneum* separately to these lineages is not possible.

##### 3.1.3. Systems of axon tracts innervating the VMC

Two tract systems, BAla and BAmv, that enter adjacent to the AL but then continue medially into the VMC, were described above. A third system, the basoanterior postero-lateral system (BAlp) approaches from the opposite side. Two tracts converge from anteriorly on to the cleft between VMC, VLP, and AMMC, and project posteriorly as the loVL fascicle (Figure 4B3, C1-3). The tracts correspond in entry and trajectory to the BAlp1-3 system defined in *D. melanogaster* (Figure 5A1-2, B1-2).

The *D. melanogaster* BAlp system includes four lineages. As just mentioned, BAlp1-3 project posterior-medially and innervate the VMC. A fourth one, BAlp4, projects anteromedially towards the posterior surface of the AL, and innervates the AL, TR and VMC. Neurons of this lineage, in particular all secondary neurons, have ascending axons that follow the antennal lobe tract and reach the superior protocerebrum. A tract clearly homologous to BAlp4 is not discernible in *T. castaneum*. However, a thick fibre bundle approaches the BAla entry point from a ventroposterior direction (which would be similar to the trajectory of BAlp4 in *D. melanogaster*). It is conceivable that this tract (termed “BAlap”; Figure 4C1) corresponds to BAlp4 (Figure 5A1, B2).

##### 3.1.4. AMMC

Aside from the posterior branch of the antennal nerve and the fibre tracts forming part of the BAla system described above, one additional fibre system enters the AMMC; we designated this as the ventral AMMC system (AMMCv). Cortex-derived fibres enter the AMMC from ventrolaterally (Figure 4C2). The vAMMC system, in particular at the early larval stage, is difficult to distinguish from the antennal sensory afferents entering the AMMC at the same location. It is clearer in the prepupa and adult brain (not shown). Based on its topology we speculate that the posterior-ventral AMMC system may correspond to *D. melanogaster* lineage BAlv (Figure 5A2, B2).

#### 3.2. Protocerebrum

##### 3.2.1. Ventrolateral protocerebrum (VLP) and posterior lateral protocerebrum (PLP)

Aside from the massive input from the optic lobe (not discussed in this work), the PLP and VLP are innervated by tract systems belonging to the basolateral (BL) groups.

(1) BL antero-ventral system (BLAv): four thick tracts enter the VLP from anteriorly. Two of these are close to each other and project straight axons posteriorly (Figure 2B1; Figure 4B4, C1-3). Upon reaching the neuropil surface, part of the fibres turn medially into the VLP; another component continues along the surface posteriorly towards the PLP. Two additional tracts originate more ventrally; one of them projects postero-dorsally upward, crossing the other two tracts laterally (Figure 2B3-4; Figure 4B3, C1-3). A second, thinner bundle projects straight posteriorly along the VLP surface (Figure 4C2). The BLAv tract system coincides with lineages BLAv1, BLAv2 and BLAvm defined in *D. melanogaster* (Figure 5A1-2, B1-2). Here, all three of these lineages have distinct hemilineages with separate tracts directed medially and posteriorly, respectively. The medial hemilineages of BLAv1 and BLAvm traverse the anterior VLP, pass underneath the peduncle, and then turn upward to form the superior arch commissure (SAC). The medial branch of BLAv2 also forms a commissural system crossing further posteriorly. Posterior components of BLAv1/BLAv2 form part of the great commissure. It is possible that *T. castaneum* counterparts connect to these commissural systems in a similar way, but global labelling of fibres with tubulin did not allow us to positively make this claim.

(2) BL ventro-anterior system (BLVa): Two bundles originating in the posterior ventral cortex, flanking the lobula complex, converge on a point located in the dorsal VLP, slightly ventrally of the LH (Figure 2B5; Figure 4B4, C1-3). Axons then continue dorsally into the LH. They might correspond to lineage tracts of the BLVa system in flies (Figure 5A2, B2). Here, two lineages (BLVa1 and a2 project dorsally, similar to the *T. castaneum* BLVa system; two additional ones (BLVa3 and a4) have short tracts terminating in the ventral VLP compartment. Tracts with these latter characteristics were not detected in *T. castaneum* (Figure 5A2, B2).

(3) BL posterior system (BLP): 7-8 axon bundles emerging from cell bodies in the posterior cortex converge and project anteriorly, passing the optic stalk (connection between optic lobe and central brain) dorsally or ventrally, and continuing along the surface of the PLP/VLP and LH (Figure 2B5-7; Figure 4B2-4, C1-2&4). *D. melanogaster* lineage tracts with these attributes belong to the BLP system. Two of these, BLP1 and BLP2, project anteriorly (similar to the *T. castaneum* “BLPv” subset of BLP in Figure 4C2); two others (BLP3 and 4) turn dorsally into the LH, similar to *T. castaneum* groups BLPi and BLPd (Figure 2B5-6)

(4) BL ventroposterior system (BLVp): 2 tracts enter slightly medially of the BLPv tracts. The dorsal one is formed by a single bundle; the thicker, ventral one by 3-4 converging bundles (Figure 2B7; Figure 4B2, C1&4). Both tracts come close at the neuropil surface; thereafter, fibres project forward as the ventral component of the PLF (PLFv; see above). The pair of *D. melanogaster* lineages BLVp1 and BLVp2, each one with a medially and an anteriorly projecting hemilineage, closely match the features of the BLVp system in *T. castaneum* (Figure 5A2-3, B2-3).

##### 3.2.2. Mushroom body, central complex and inferior protocerebrum

*Mushroom body:* Neurons of the MB are located in the dorsoposterior cortex, and belong to two lineages (Zhao et al., 2008), referred to as MBm and MBl, respectively (Figure 4E1-3). Accordingly, the calyx (CA) of the MB, which receives dendritic arborizations close to the neuronal cell bodies, is composed of two lobes, a medial and a lateral lobe, visible from early larval stages onward (Figure 4E1-3). Axons of MB neurons assemble in a large number of individual tracts that enter the calyx from dorsoposteriorly, forming two massive roots (Figure 2B4-5, D5; Figure 4B5-6, E1-3). Both roots fuse into the peduncle (PED), which projects forward all the way to the anterolateral brain surface (Figure 4E1).

*Central complex:* The vast majority of neurons of the CX are the columnar and pontine neurons whose fibres enter from posteriorly in close association with the protocerebral bridge (PB) and then continue forward into FB, NO and EB. These neurons are formed by four type 2 lineages, DM1-DM4, which due to their peculiar proliferation pattern (Bello et al., 2008; Boone & Doe, 2008; Bowman et al., 2008) are much larger than “regular” (type 1) lineages and form multiple, closely spaced tracts, rather than one or two. The topology of these type 2 lineages appears to be highly conserved among insect taxa (Boyan et al., 2017; Farnworth et al., 2020). DM1-3 form the dorsoposterior medial group of tracts (DPMm/pm); DM4 belongs to the central medial group (CM). DM1 is located furthest medially/anteriorly, and DM4 most laterally/posteriorly. Note that only part of the neurons generated by DM1-4 target the CX; other neurons of these lineages also innervate other compartments.

In the *T. castaneum* larva, each type 2 lineage has 4-6 bundles that converge onto three fascicles, the roots of the fan-shaped body. They comprise the lateral root (lrFB; tract of DM4), dorso-lateral root (dlrFB; DM3 and DM2;) and medial root (mrFB; DM1; Figure 2B5-6, D5; Figure 4B4-5, E1-2; Figure 5A3, B3). Reaching the FB, the roots split up into the posterior plexus of the fan-shaped body (pplFB; Figure 4B4). DM4 axons destined for the lateral root initially extend along the medial surface of the medial equatorial fascicle (MEF) before turning sharply medially into the pplFB (Figure 4B4, E1-2). The dlrFB has a unique relationship to the medial antennal lobe tract (mALT) that intersects this root at a right angle (Figure 2B4, D4; Figure 4B4, E1). As described for *D. melanogaster*, the DM3 contingent of fibres passes the mALT laterally, the DM2 contingent medially (Figure 2B4, D4). The mrFB enters the plexus from straight posterodorsally (Figure 2B4, D4; Figure 4B4, E1). A GFP-tagged marker of the gene Rx, expressed in subsets of columnar neurons of DM1-DM4, has revealed in a recent study (Farnworth et al., 2020) that the pattern in which these neurons decussate and contribute to the columns of the FB and EB also coincides with the canonical pattern known for flies and locusts (although the relative timing differs substantially).

*Central domain of the inferior protocerebrum (IPc):* A large number of tracts formed by neurons located in the posterior brain cortex project onto the IPc at its borders with the SMP/SLP and VMC/PLP from posteriorly. We first consider tracts entering dorsally and medially of the mushroom body (the CM and DPM super-sets of tracts), and subsequently those located laterally of this structure (CP). Among the medial tracts one can distinguish:

(1) The CM dorsal system (CMd): This large tract system is formed by two smaller “subsystems” of axon bundles that consists of 4-6 converging tracts each.

(1.1) The medial component of the CMd system belongs to DM4, mentioned in the previous section (Figure 2B6, D2; Figure 4B3-4, D1-3). Axons project straight anteriorly into the MEF; part of them leave the MEF medially as the lrFB (see above). Most axons continue anteriorly along the MEF before turning medially as the LALC commissure.

(1.2) The lateral component of CMd, formed by at least 5 tracts, is located laterally adjacent to DM4. Projecting forward it forms a distinct lateral bundle within the MEF (Figure 2B6, D2; Figure 4B3-4, D1-3). Reaching the level of the great commissure (GC), this bundle leaves the MEF and turns ventromedially into the GC (MEF to GC, Figure 2B4, D2). An axon tract with these characteristics is typical for *D. melanogaster* lineage DPMpl3; what stands out in *T. castaneum* is the much larger number of neurons (and individual tracts), as well as the closer proximity to DM4 (Figure 5A3, B3).

(2) The CM ventral system (CMv): Ventrally of DM4 and DPMpl3 we find two other groups of tracts, one more medially, the other more laterally (Figure 2B6-7; Figure 4B2, D2-3). Together, they correspond in position and projection to *D. melanogaster* type 2 lineages DM5 and DM6; we annotate them as DM5/6m and DM5/6l. Most fibres of DM5/6 follow along the MEF anteriorly; some then turn ventrally towards the GC, similar in trajectory to DM6 in *D. melanogaster*. In addition, ventrally directed branches of DM5/6m form the longitudinal ventroposterior fascicle (loVP; Figure 2D5; Figure 4D2-3).

(3) The CM medial system (CMm): consists of 2-3 thin tracts that are formed by neurons located medially of DM4, and that converge onto the medial entry point of the MEF (Figure 4B3, D3. In location, they resemble *D. melanogaster* lineage CM5.

(4) The DPM posterolateral system (DPMpl): Two converging tracts enter at a position dorsally of DM4/DPMpl3 (Figure 2B4-5, D2-4; Figure 4B5-6, E1-3). Instead of joining the MEF, they turn dorsally and form the loSM. Their axons project straight anteriorly, skirting the adjacent medial root of the calyx (Figure 2B5) and continuing forward to a level in front of the central complex. *D. melanogaster* DPMpl1/2 exhibit the same characteristics as the *T. castaneum* DPMpl tracts (Figure 5A3, B3). In addition to the DPMpl tracts, branches of DM3 and DM4 can be also observed (just as in *D. melanogaster*) to contribute to the loSM fascicle.

As outlined in the previous section, the lateral neuropil of the posterior hemisphere is dominated by three fascicles which demarcate the border between IPc and adjacent neuropils, the lateral equatorial fascicle (LEF), posterior lateral fascicle (PLF), and oblique posterior fascicle (obP). In *D. melanogaster* the four lineages of the central posterior group (CP1-4) and three lineages of the dorsoposterior lateral group (DPLp1/2, DPLpv), form the fibres contributing to these fascicles. We distinguish in the *T. castaneum* larva 10-12 fibre tracts with general attributes of *D. melanogaster* CP1-4, DPLp1/2, and DPLpv. In all, they give rise to neuropil fascicles (LEF, PLF, obP) resembling those of their *D. melanogaster* counterparts. However, the ratio of fibres contributing to these bundles differs: much less fibres form part of the obP and PLF, and more contribute to the LEF. In addition, a fascicle altogether missing in flies, the ventral obP (obPv), has a strong presence in *T. castaneum*.

(1) The CP dorsal system (CPd): this fibre system consists of four thin tracts that converge at the neuropil boundary between IPc and calyx, and then turn dorso-medially over the peduncle as the obPd fascicle (Figure 2B5; D4-5; Figure 4B5-6, D1-3).

(2) The DPL posterior system (DPLp): A second component of the obPd is contributed by part of the fibres of a pair of converging bundles coming from more laterally (DPLp; Figure 2B5-6; Figure 4B5-6, D1-3). Other fibres of this tract system diverge to project anteriorly, which matches the trajectories exhibited by *D. melanogaster* lineages DPLp1/2 (Figure 5A2-3, B2-3).

(3) The CP ventral system (CPv): two tracts converge onto the boundary between IPc and ventrally adjacent VCM, and project forward as the lateral equatorial fascicle (LEF; Figure 2B6-7, D5; Figure 4B3, D1-3). Lineage CP1 forms the LEF in *D. melanogaster* (Figure 5A2-3, B2-3)

(4) The CP intermediate system (CPi): Four tracts dorsally adjacent to the CPv pair, annotated as CPi, converge to then contribute to two neuropil fascicles. The first one joins the LEF; the second one bends dorso-medially and passes the peduncle at its ventral surface (Figure 2B6, D5; Figure 4B3, D2-3). A comparable fascicle does not exist in flies; we will call it the ventral obP (obPv), because it extends parallel to the dorsal (“canonical”) obP (which passes dorsally of the peduncle), and, like the obP, turns into the loSM system (Figure 4B4).

(5) The CP/DPL posteroventral system (CP/DPLpv): A system of 3-4 tracts converge on the boundary between IPc and PLP and, continuing forward, appear to give rise to both PLFdm and PLFdl (Figure 2B5, D1&5; Figure 4B3, D1-3). In *D. melanogaster*, the ventral hemilineages of CP2 and 3 (CP2/3v) form PLFdm; lineage DPLpv forms PLFdl (Figure 5A2-3, B2-3).

*Posterior inferior protocerebrum (IPp):* The IPp, located in between the protocerebral bridge and IPc, is contacted by the tract systems that form the roots of the fan-shaped body, as well as those collecting into the ME F and loSM. These were described in the previous sections. In addition, the IPp is approached at its dorsal surface by the tract system we here call DPM dorsal (DPMd). Two tracts converge at the neuropil surface and form a thick bundle that extends ventrally, passing the mALT/dlrFB laterally, before it is lost in the neuropil of the VMC (Figure 2B4; Figure 4B6, E1-3). The third tract, located more medially, also extends vertically downward, and is lost in the IPp neuropil. Regarding location and vertical trajectory, these bundles are likely to correspond to the DPMl1/DPMl2 lineages (lateral tract) and the DPMm2 lineage (medial tract) of *D. melanogaster* (Figure 5A3, B3). DPMl1/2 generate one of the major descending neuron populations whose axons form the dorsoposterior protocerebral tract (DPPT; Lovick et al., 2013); global labelling with tubulin shows the beginning of a descending bundle in *T. castaneum*, but this bundle cannot be followed far (Figure 4E1, 2).

*Anterior domain of interior protocerebrum (IPa) and LAL:* A number of tract systems formed by neurons located in the anterior and antero-lateral cortex enter the IPa and ventrally adjacent LAL. These tracts belong mainly to the DAL super-set.

(1) The DAL centrolateral system (DALcl): two fibre bundles formed by cell bodies located laterally of the IPa converge and enter the neuropil in the shallow groove between IPa and LAL (Figure 2B1; Figure 4B6, F1-3). They correspond in entry point and further trajectory to the lineage pair DALcl1/DALcl2 in *D. melanogaster* (Figure 5A1-2, B1-2). Thus, similar to DALcl1/DALcl2, the *T. castaneum* al IP system bifurcates into a ventral and dorsal branch. The ventral branch projects postero-medially, defining the border between LAL and IPa. The branch then reaches the mushroom body medial lobe, passes under its ventral surface, and then contacts the central complex. Part of the fibre bundle seems to enter the lateral surface of the central complex as the anterior lateral ellipsoid tract (LEa; better visible in adult); most fibres turn medially and cross the midline in the subellipsoid commissure (SuEC; Figure 2B2; Figure 4B6; F1&3). The dorsal branch of the DALcl system follows a posterior trajectory, resembling the dorsal hemilineage tract of *D. melanogaster* DALcl1/2. It passes the vertical lobe of the mushroom body at its lateral side, and then turns medially and ventrally, joining a thicker bundle of axons formed by the DALcm system (Figure 4F1-2; see below). The resulting composite tract corresponds to the descending central protocerebral tract (deCP).

(2) The DAL centromedial system (DALcm): A pair of tracts converges at the dorsolateral IPa surface. It splits up into a medial branch that continues posteromedially into the IPa, towards the ML, and a lateral branch that continues posteriorly, then curving medially and ventrally around the posterior surface of the VL (Figure 2B1, C1, D1&3; Figure 4B7, F1&3). Joined by DALcl (see above) the tract continues ventrally, forming the conspicuous deCP tract (Figure 4B5-6, F1-3). In *D. melanogaster*, lineage pairs DALcm1/DALcm2 resemble the DALcm system, forming a medial tract (DALcm1/2m) into the IPa (Figure 5A1, B1), and a ventral tract (DALcm1/2v) into the deCP. A third lineage, DALd (with axons descending into the VNC) joins DALcm1/2v in the deCP (Figure 5A1). It is unclear whether such descending DALd fibres exist in *T. castaneum*, possibly forming an integral part of the two tracts of the DALcm system.

(3) The BA mediodorsal system (BAmd) is formed by two tracts. One, located at a more lateral position, has a highly conspicuous branch point that occurs prior to entering the neuropil. The dorsal branch converges on the DALcm tract and continues into the IPa; the ventral branch projects ventroposteriorly, following the anterior surface of the IPa, then turns medially and joins a commissural fibre bundle that crosses the midline in the ALC (Figure 2B1; Figure 4B5-6, C3). The second BAmd tract lies more medially but follows the same course as the ventral branch of the lateral tract; turning medially, it also crosses ALC (Figure 2B2; Figure 4C3). Two lineage associated tracts with these exact features are represented by BAmd1d/v and BAmd2 in *D. melanogaster* (Figure 5A1, B1). Here, the commissural bundle formed by these lineages is also part of the antennal lobe commissure. Adult flies possess a massive ALC, predominantly composed of crossing olfactory afferents. Such crossed sensory afferents are missing in the *D. melanogaster* larva, and the ALC is exclusively formed by the BAmd1/BAmd2 lineage pair. In the *T. castaneum* larva, crossing fibres other than BAmd seem to exist (Figure 2B2); these are possibly crossed olfactory afferents.

(4) The DAL ventrolateral system (DALvl): two fibre bundles enter the depression in-between the IPa, LAL and VLP (Figure 2B1; Figure 4B5-6, F1-2). The ventral one of these tracts shows a highly characteristic trajectory which clearly identifies it with the *D. melanogaster* DALv1 lineage tract (Figure 5A1-2, B1-2). Thus, the bundle (called LEFa) extends posteriorly along the lateral surface of the LAL. It runs more or less parallel to the peduncle of the mushroom body, helping to define the ventral boundary of the IPc compartment (Figure 2B2-3); axons of the DALv1 tract then turn ventrally and radiate into the great commissure. The second component of the DALvl system enters dorsally of DALv1 (Figure 4B6, F1-3), but soon converges with it, at which point it cannot be separately followed. Based on position and closeness to DALv1 this bundle could correspond to *D. melanogaster* lineage tract DALl1 (Figure 5A1-2, B1-2).

(5) The DAL ventromedial system (DALvm): This system is comprised of three separate tracts. One follows the lateral LAL surface posteriorly, giving off fibres into the LAL, but apparently mainly targeting the VMC posteriorly adjacent to the LAL (Figure 2B1; Figure 4B5, F1-2). Two additional tracts enter the LAL at its dorsal and ventral boundaries, respectively, from anteriorly (Figure 2B1-2; Figure 4B5-6, F1-3). The DALvm system becomes more prominent in the adult brain, where it converges with the ventral branch of the DALcl1/2, projecting underneath the ML towards the central complex and contributing to the LEa fascicle (see above). In terms of entry point and relation to DALcl, the DALvm system resembles closely the lineages DALv2/DALv3 in *D. melanogaster* (Figure 5A1, B1).

##### 3.2.3. Superior protocerebrum

*Superior medial protocerebrum:* Several tract systems (e.g., DPMpl) that enter the neuropil at the SMP/IPc/IPp boundary from posteriorly were described in a previous section. Additional tract systems entering the SMP from anteriorly and laterally belong to the DAM and DPL groups of tracts, respectively.

(1) The DAM ventral system (DAMv): Two tracts approaching from straight anteriorly converge in the cortex, resulting in a thick bundle that continues into the SMP. Part of the fibres appear to turn medially into the supraesophageal commissural system, forming the most anterior and dorsal component of this system. Based on location and trajectory, the anteromedial SMP system corresponds to *D. melanogaster* lineage tracts DAMv1/DAMv2; the commissure represents the anterior-dorsal commissure (ADC; Figure 2B1; Figure 4B6, F1-3; Figure 5A1, B1).

(2) The DAM dorsal system (DAMd): 3-4 tracts converge within the anterior dorsal cortex, medially of the mushroom body VL (Figure 2B2; Figure 4B7, F1-3), to form a thick bundle; reaching the anterior SMP surface, fibres turn posteriorly, projecting towards the loSM fascicle, and therefore called loSMa (Figure 2B2; Figure 4F2-3); other fibres continue ventromedially into the ADC commissure. In regard to these properties the anterior dorsal system corresponds to the tracts of the DAMd1-3 lineages in *D. melanogaster* (Figure 5A1, B1)

(3) The DPL central system (DPLc): this is a complex group of tracts entering the neuropil vertically at the boundary between SMP and SLP. We distinguish a medial component and a lateral component. The medial component (DPLcm) has four tracts that converge at the neuropil surface, two entering from dorsoanteriorly, with somata laterally adjacent to the VL (Figure 2B3; Figure 4B7, F1-4), and the other two entering from dorsoposteriorly. These tracts converge upon the trSPm fascicle (see above). The medial tract of the posterior pair bifurcates at neuropil entry, with one branch extending posteroventrally. This bifurcated tract resembles DPLc5 in *D. melanogaster*; the other three topologically represent DPLc3 (the posterior lateral tract) and DPLc1/DPLc2 (anterior pair; Figure 5A1-2, B1-2).

The lateral component of the DPLc system (DPLcl), resembling the *D. melanogaster* lineage tract DPLc4, and consisting of two thick converging tracts, enters slightly laterally of the medial component (Figure 2B3; Figure 4B7, F1-4). Its fibres also turn medially, joining the trSPm.

(4) The DPL medial system (DPLm) is represented by 2-3 converging bundles entering the neuropil at the junction between calyx (posterior-medially), SMP (anterior-medially), and SLP (laterally; Figure 2B4; Figure 4B6, F1-4). Fibres of this system may likely correspond to the *D. melanogaster* DPLm1/DPLm2 lineages (Figure 4A2-3, B2-3).

*Superior lateral protocerebrum (SLP) and lateral horn (LH)*: Tracts entering the superior protocerebrum from lateral belong to the DPL super-set. They include the following four systems.

(1) The DPL anterior system (DPLa): 3-4 bundles converge at the border between SLP (medially), VLP (ventrally) and LH (posteriorly; Figure 2B3; Figure 4B7, F1-3). Most of their fibres give rise to the trSA fascicle, which in *D. melanogaster* is formed by lineages DPLal1-3 (Figure 5A1-2, B1-2). A fourth *D. melanogaster* lineage of this system, DPLam, lies dorsomedially of DPLal1-3 and projects straight through the virtual centre of the trSA arch. A similar configuration is clearly visible in the adult *T. castaneum* brain, and faintly evident in the larva (Figure 2B2-3).

(2) The DPL posterolateral system (DPLpl): 4-5 converging fibre bundles contact the lateral surface of the SLP at a level posteriorly adjacent of that of DPLal/DPLam (Figure 2B4; Figure 4B5-6, F1-2&4). In the larva they are very close to DPLa; in the prepupa and adult, the growth of the SLP has created a considerable distance between the DPLa and DPLpl systems (Figure 7). Part of the fibres project anteriorly at the SLP surface towards the entry point of DPLa; however, most fibres enter the neuropil at the border between SLP (medially) and LH (ventrolaterally) and follow an anterior ventral course that brings them close to the peduncle (Figure 2B4; Figure 4B5-6, F1). The DPLpl system shares topological features with the *D. melanogaster* lineage pair DPLl2/DPLl3 (Figure 5A2, B2). However, the trajectory of fibres of the *T. castaneum* DPLpl system within the neuropil is significantly different from that of the *D. melanogaster* DPLl2/3. Here, the deep component (hemilineages) of this lineage pair projects straight anteriorly, forming the longitudinal superior lateral fascicle (loSL), as opposed to *T. castaneum*, where axons project ventromedially towards the peduncle (Figure 2B4, D4-5; Figure 4B6-7, F1)

(3) DPL posteromedial system (DPLpm): A thick bundle reaches the SLP at a point located posteriorly and medially of DPLpl (Figure 2B4; Figure 4B6, F1-2&4). It enters the neuropil and projects anteromedially, coming close to the loose trSPm fascicle formed by the DPLc system of tracts (Figure 3F1). A lineage projecting from a posterolateral position towards the DPLc system in *D. melanogaster* is DPLl1; what is different to the observed pattern in *T. castaneum* is that DPLl1 enters in close proximity to DPLl2 and DPLl3, rather than choosing its separate entry portal (Figure 5A1, B1).

(4) The BL anterodorsal system (BLA/D): 5-6 fibre bundles converge upon the neuropil surface at the junction between SLP (anterior), LH (posterior) and PLP (ventral). Part of the bundles approach this entry point from posteriorly, others from anteriorly. The bundles pass right underneath the anterior optic tract and then loop upward to form the trSI fascicle (Figure 2B3-4; Figure 4B5-6, F1-4). Fibre systems with all of these attributes are lineages of the BLD group (BLD1-4; posterior) and the BLAd group (BLAd1-4; anterior; Figure 4A2, B2).

### 4. Comparison of patterns in *T. castaneum* and *D. melanogaster* brain development

We wanted to get a rough estimate of brain sizes in *T. castaneum* and *D. melanogaster* at different developmental stages. For this, we did simple measurements along the major geometric axes and calculated an ellipsoid volume from these measurements to get a growth estimation from one stage to another (Table 4). We found the highest similarity in volume at the late larval stage between the species, and a very pronounced increase during the larval stage, and a much-reduced increase in volume during pupal phases. Note that in *D. melanogaster* we used L3 (wandering third instar larva) brains and not prepupae, as in *T. castaneum*, so there might be some larval growth included from L3 to adult.

**Table 4:** Brain size measurements along the three geometric axes as well as a calculated ellipsoid volume, to illustrate growth during larval and pupal stages and growth similarities between, both, stages and species. Raw sizes are in µm. Indicated are means and the standard deviation from individuals from each species. % increase indicates the amount of change from L1 to end larval stage and from end larval stage to adult in ellipsoid volume.

| Species | Stage (N) | Width | Height | Depth | Volume ($\times 10^6$) | % increase |
| --- | --- | --- | --- | --- | --- | --- |
| *T. castaneum* | L1 (11) | 141.13 ± 23.28 | 133.02 ± 21.05 | 87.36 ± 12.55 | **7.09** ± 2.50 | - |
| | Prepupa (10) | 377.54 ± 52.47 | 291.68 ± 33.10 | 139.49 ± 37.72 | **64.26** ± 23.00 | 907 |
| | Adult (12) | 559.32 ± 37.94 | 311.27 ± 14.45 | 107.57 ± 33.55 | **77.43** ± 20.36 | 120 |
| *D. melanogaster* | L1 (5) | 99.28 ± 4.45 | 60.20 ± 9.63 | 57.60 ± 3.05 | **1.45** ± 0.32 | - |
| | L3 (5) | 369.60 ± 32.57 | 187.00 ± 24.71 | 175.00 ± 17.38 | **51.02** ± 11.22 | 3511 |
| | Adult (5) | 594.40 ± 27.51 | 296.20 ± 9.47 | 203.80 ± 21.43 | **149.78** ± 11.59 | 294 |

In both species, brain growth results from an increase in cell number and individual cell size. A comprehensive study of the dynamics of neuroblast proliferation and lineage development for *T. castaneum* was out of the scope of this work, but several differences in these aspects set the beetle apart from the fly, and these will be highlighted in the following section.

Active neuroblasts can be recognized by their size, superficial position and high mitotic rate from the embryonic stages onwards when they first delaminate from the ectoderm. Neuroblasts, amounting to approximately 100 in number, feature prominently in both *T. castaneum* and *D. melanogaster* during early development (Urbach et al., 2003; Urbach & Technau, 2003a). At that stage, neuroblasts and their progeny, including ganglion mother cells (GMCs) and postmitotic neurons with their outgrowing axons form conspicuous, evenly spaced packages that make up the embryonic brain primordium. At late embryonic stages, neuroblasts in *D. melanogaster* cease to proliferate for about 24h, and as a result, in the freshly hatched larva, almost no neuroblasts are visible (Figure 6A), except for the 4 neuroblasts of the mushroom body, and one neuroblast associated with the antennal lobe. Beginning at the L1-L2 transition, neuroblasts enlarge and start dividing again, and at the late larval stage, a configuration resembling the mid-stage embryo reappears, with large neuroblasts at the surface, followed by GMCs and newly produced secondary neurons forming axon tracts at the surface of the brain cortex (Figure 6B). Differentiated primary neurons produced in the embryo are distinguished from their younger siblings by their large cell size and deep location next to the neuropil (Figure 6C). The secondary neurons do not differentiate immediately. Their axons form bundles that invade the neuropil (alongside the primary axon tracts; Larsen et al., 2009) and, augmented in volume by tufts of filopodia, form large “lacunae” devoid of synapses. These lacunae, which clearly stand out in brain preparations labelled with synapse markers (e.g., Brp; Figure 6E), are visible through most brain regions, notably the central complex, antennal lobe, ventromedial cerebrum and superior lateral protocerebrum (Lovick et al., 2017). Differentiation of secondary neurons takes place during the pupal period. As a result, the neuropil grows in volume, while, at the same time, the cortex, formed by cell bodies that no longer increase in number, thins out. These changes in the cortex also affect the appearance of lineages. Thus, neurons related by lineage, forming a narrow cylinder in the late larval brain cortex, spread out horizontally in the thinning cortex; the axon tracts associated with each lineage also widen, and become less easily defined compared to the larval stage (Figure 6D).

**Figure 6:**
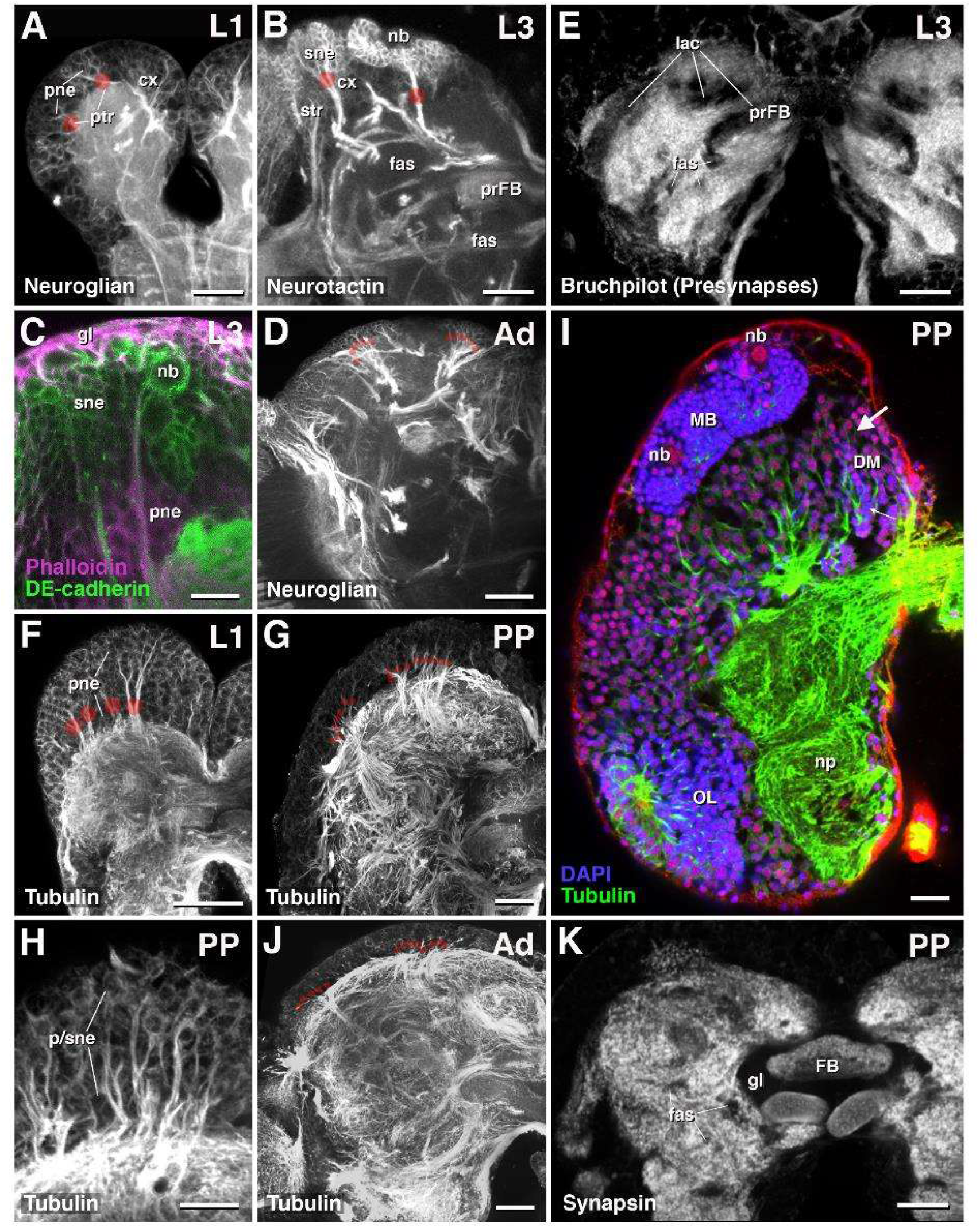
Features of neural development in the larva of *D. melanogaster* and *T. castaneum.* All panels show z-projections of frontal confocal sections of brains at different developmental stages (L1: early larva; L3: late larva; PP: prepupa; Ad: adult) labeled with global neural markers as indicated at panel bottoms. (A-E): *D. melanogaster*; (F-K): *T. castaneum*. (A) Primary neurons (pne) and their tracts (ptr) form the early *D. melanogaster* larval brain. (B, C) Secondary neurons (sne; marked by Neurotactin and DE-cadherin) form dense clusters of small cells, proliferating from neuroblasts (nb) located at the brain surface. They project secondary axon tracts (str) into the neuropil. These tracts appear as distinct, cohesive bundles, in the cortex and at neuropil entry (large red dots). (D) Secondary axon tracts in the adult fly brain cortex, which is much thinner than the larval cortex, form frayed, wider bundles (small red dots). (E) Late *D. melanogaster* larval brain exhibits multiple large, synapsin-negative lacunae (lac), which are formed by undifferentiated secondary axons and filopodia. (F) Early larval brain of *T. castaneum* with primary neurons and axon tracts. (G) The thinning of cortex and widening/fraying of axon tracts (small red dots), which happens during metamorphosis in *D. melanogaster*, occurs already during larval stages in *T. castaneum*. (H, I) Large, superficially located neuroblasts cannot be recognized in the late *T. castaneum* larval brain, with the exception the two mushroom body neuroblasts. Furthermore, the distinct zonation into a deep layer of large primary neurons and a superficial layer of small secondary neuron typical for the late *D. melanogaster* larva (see panel C) is not apparent in *T. castaneum*. (K) No synapsin-negative lacunae can be detected in the larval *T. castaneum* brain. For all abbreviations see Table 2. Scale bars: 25µm (all panels except C, H); 10µm (C, H).

**Figure 7:**
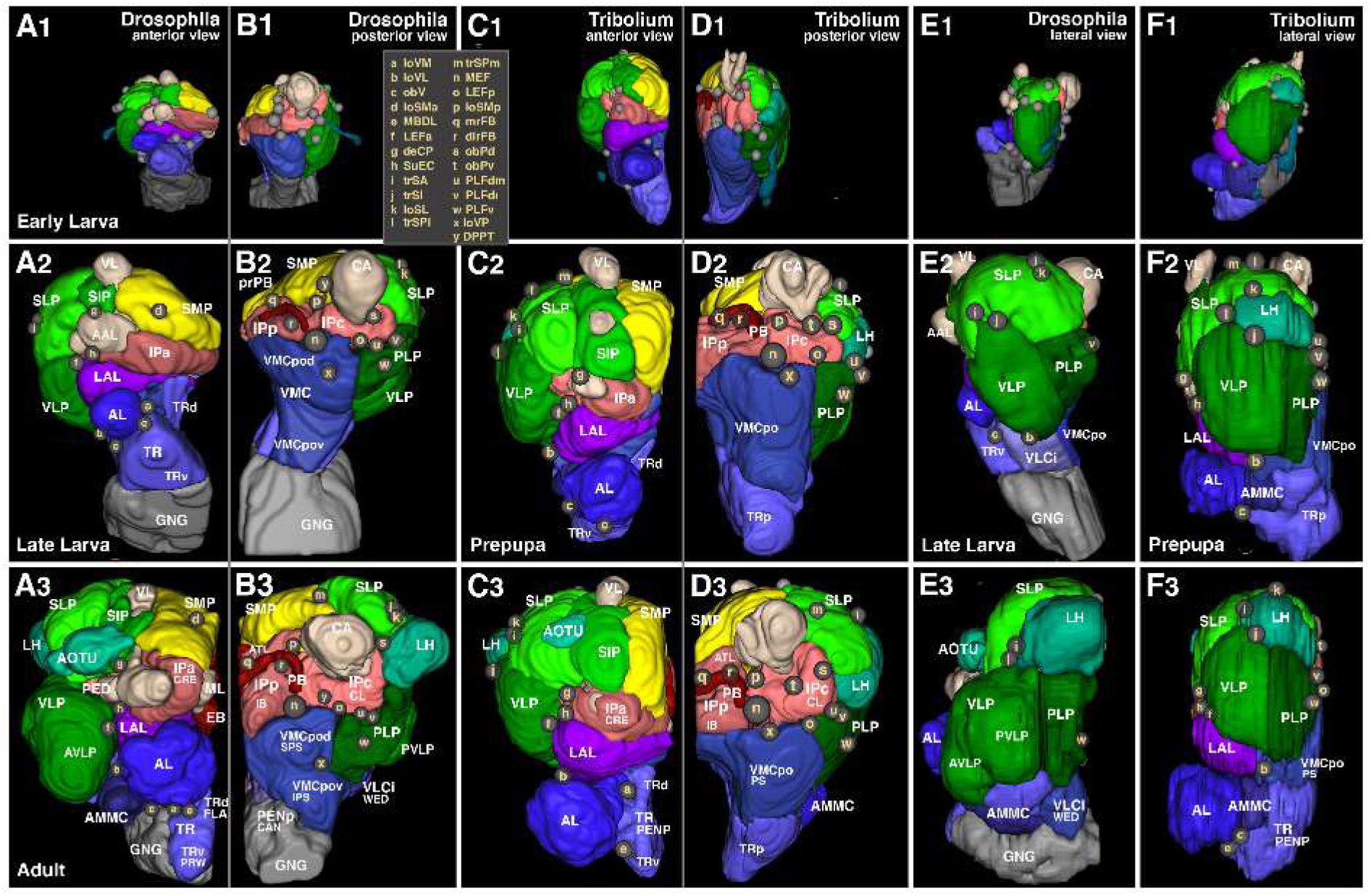
Comparative growth of brain compartments in *D. melanogaster* and *T. castaneum*. Panels show digital 3D models of one brain hemisphere of early larva (top row), late larva/prepupa (middle row) and adult (bottom row) in anterior (A1-3, C1-3), posterior (B1-3, D1-3) and lateral view (E1-3, F1-3), all to scale. Neuropil compartments are rendered in the same colors as in previous figures (Figures 2–5) and are annotated in white lettering. As in previous figures (Figs.2–4), small spheres annotated in beige single letters represent locations where neuropil fascicles enter the neuropil. For correspondence of single letters with fascicle names see inset in B1 and C1. For all abbreviations see Table 2. *The D. melanogaster* larval models are based on Cardona et al. (2010) and the adult model is based on Pereanu et al. (2010).

The brain cortex of the freshly hatched *T. castaneum* larva resembles that of *D. melanogaster*. Primary neurons form packages, emitting discrete axon bundles, as described in the previous section (Figure 6F). Neuroblasts (with the exception of two large neuroblast associated with the mushroom body) are no longer visible. However, during the ensuing larval period, lineage development appears to differ between the two species. At the end of larval life, with the exception of MB neuroblasts, *T. castaneum* still shows no large, neuroblast-typic cells in in the brain cortex (Figure 6G). At mid-larval stages, only few scattered large cells are visible at the brain surface (not shown). Furthermore, the characteristic arrangement of small-diameter secondary neurons at the surface and large-sized primary neurons in the depth of the cortex is not apparent (Figure 6H, J). Large clusters of small-sized secondary neurons are evident in the vicinity of the mushroom body neuroblasts, and at a few other locations (notably the tritocerebrum, and the optic lobe; Figure 6J); however, in most domains of the cortex, large and small cells appear intermingled, and also do not appear as two discrete classes as in *D. melanogaster*. Large-sized neurons are equally likely to appear at the cortex surface or near the neuropil (Figure 6H, J). The tendency of axon tracts to spread out into more loosely textured bundles, a process that occurs during metamorphosis in *D. melanogaster*, is already evident in the late larva or prepupa of *T. castaneum* (Figure 6G), which thereby resembles the adult brain (Figure 6I). Both, the patterns of active neuroblasts and the timing of cortex spreading indicate that the second proliferation phase occurs earlier in *T. castaneum* than in *D. melanogaster*.

A further difference between the two species that may be related to the proliferation and differentiation of neurons concerns the structure of the late larval neuropil. In late *D. melanogaster* larvae, voluminous lacunae filled by axons and filopodia of undifferentiated secondary neurons occur in the neuropil. Such domains are absent in the late *T. castaneum* larval brain. This has been shown in detail for the central complex, which contains many differentiated neurons already in the larva (Farnworth et al., 2020), whereas it constitutes a non-functional, asynaptic primordium in the fly (Andrade et al., 2019; Lovick et al., 2017). However, it appears that the same difference applies to the brain in general. Thus, aside from the synapse-free channels filled by fascicles, or the massive synapse-free glial layer surrounding the central complex, no other volumes lacking synapses are detectable (Figure 6K). This indicates that the connectivity of secondary neurons develops with different timing in the two species: delayed in the fly but soon after a neuron's birth in the beetle.

When looking at the growth of different parts of the brain in *T. castaneum* in comparison to the fly, we see many conserved features. Compartments with strongest growth are first and foremost the sensory centres, antennal lobe (Figure 7A1-3, C1-3) and optic lobe (not shown). This results from the strong increase in the number of the corresponding sensory neurons and their terminals. An interesting difference in growth dynamics relates to the AMMC, the sensory domain receiving mechanosensory input from the antenna. In *D. melanogaster*, the AMMC and its innervating elements are mainly secondary, generated in the late larva and pupa. Thus, in the fly larva (with the possible exception of a single sensory axon; see Kendroud et al., 2017), no antennal nerve fibres pass beyond the AL, and an AMMC is still missing in early and late larvae (Figure 7E1-3). By contrast, in *T. castaneum*, a rudimentary AMMC (innervated by distinct AN branch and BAlc fibres; see section above) exists already in L1 (Figure 7F1-3). Similarly, the PB and FB emerges in the *T. castaneum* embryo but in the *D. melanogaster* pupa (Farnworth et al., 2020).

In addition to the aforementioned sensory domains, all other compartments increase throughout postembryonic development of fly and beetle. However, the superior protocerebrum, in particular the SLP and LH, as well as the VLP, increase in volume more strongly in both species than other domains (Figure 7A-D). Growth in the VLP and LH is more pronounced in *D. melanogaster* than in *T. castaneum* (Figure 7A2-3, B2-3, C2-3, D2-3), which is likely related to the larger size of the adult optic lobe in *D. melanogaster*, since much of the output of the optic lobe is directed at the VLP (Strausfeld, 1976). The difference in growth of the anterior optic tubercle (AOTU), which also develops to a much greater size in *D. melanogaster* (Figure 7A3, C3), points in the same direction. Hence, it appears that the more developed visual sensory system in *D. melanogaster* is reflected in increased postembryonic development. A careful analysis of the formation of the optic lobe and the connections with the central brain, which was not attempted in our work, may be able to further substantiate of these growth patterns.

Finally, a fundamentally different structure of the tritocerebrum / subesophageal zone (SEZ) in *T. castaneum* and *D. melanogaster* accounts for a number of prominent growth features that distinguishes the two species. In flies, the SEZ, forming a merger of tritocerebrum and the three gnathal neuromeres, becomes closely connected to the supraesophageal ganglion along the entire antero-posterior axis during metamorphosis loosing clear morphological separation (Figure 7B2-3, E2-3). Neuronal processes derived from gnathal neurons extend upward into the supraesophageal ganglion, and vice versa (Hartenstein et al., 2018), resulting in the formation of neuropil domains that are of mixed sub- and supraesophageal origin. Notable among these are the inferior ventrolateral cerebrum (VLCi; “wedge”; Ito et al., 2014; Figure 7E3) and part of the VMC called “cantle” (CAN; Ito et al., 2014; Figure 7B3). The tritocerebrum remains clearly separated from the gnathal ganglion and both VLCi and CAN are conspicuously absent in *T. castaneum* (Figure 7D3, F3) indicating that these domains may have evolved by the merger of the SEZ with the supraesophageal ganglion.

## Discussion

### Tracts, fascicles and compartments as conserved building blocks of the insect brain

Our analysis demonstrates that it is possible to identify homologous fascicles and axon tracts with conserved projection patterns in *T. castaneum* and *D. melanogaster*. For fascicles, this was possible for all stages investigated. Axon tracts, however, stand out only at a time in development where the cortex is still thick, with lineages forming long, cylindrical units spanning the distance from cortex to neuropil. This means that the assignment of corresponding tracts between *T. castaneum* and *D. melanogaster* is restricted to certain larval stages. This stage corresponds to the third instar in *D. melanogaster*, but the first instar in *T. castaneum*, because, as cells distribute more widely, a considerable thinning of the cortex has already occurred at later larval stages in the beetle. The discrepancy may reflect the earlier and more continuous proliferation pattern in *T. castaneum* compared to the more compressed time of proliferation in the fly (see below).

As expected from previous anatomical studies (Immonen et al., 2017) we found considerable conservation in overall fascicle and compartment anatomy between *T. castaneum* and *D. melanogaster*, two species that shared a last common ancestor about 300 million years ago (Misof et al., 2014). We describe a set of anatomical characters for a large majority of axon tracts that are shared between *T. castaneum* and *D. melanogaster*. This anatomical complexity makes it highly likely that we are dealing with homologous entities, even in the absence of specific genetic markers. Among the examples detailed in the Results are BAmv3 and BAmv1/2, the BAla system, or the DALcl and DALcm systems. In some cases, genetic markers to define genetic neural lineages (Koniszewski et al., 2016), affirm the homology between tract systems, as shown recently for the marker Rx (Farnworth et al., 2020).

Tract systems and fascicles establish a conserved scaffold of anatomical features that has many more elements than the main neuropil compartments (e.g. MB, CX, etc). As a result, our atlas will make it possible to assign specific neurons (visualized by expression of specific molecules like transmitters, receptors, or regulatory genes, among others) to the identified tracts and fascicles. That is important when working within the same species, but also when comparing different species. For example, analyses of the central complex and its input pathways have agreed on homologies between relevant neuron types between multiple species, including *D. melanogaster*, honey bee, monarch butterfly, desert ant, dung beetle, and locust (Honkanen et al., 2019). Neurons carrying visual input from the optic lobe to the central complex (TuBu in *D. melanogaster*: Omoto et al., 2017; TuLAL in other insects: el Jundi et al., 2014) can be homologized based on their tract/fascicle: These neurons enter the neuropil at the convergence of the two mushroom body lobes, cross the peduncle, and terminate laterally adjacent to the ellipsoid body (a region called bulb); here they synapse onto another class of neurons (R-neurons in *D. melanogaster*; TL2 and TL3 in other species) which can also be homologized based on a highly conserved tract pattern. Hence, when combining a global marker that visualizes tracts and fascicles, such as Tubulin, with specific markers for cell types it should be possible to extend the identification of neurons and their homologization across species boundaries for most cells of the brain, beyond selected key examples such as the highlighted visual pathway cells.

### Differences within the conserved pattern

Despite the overall similarity in the anatomy of tracts, fascicles and compartments between *T. castaneum* and *D. melanogaster*, differences do exist. Some of the most obvious concern the general size and shape of compartments, such as the optic lobe and its targets, the AOTU, VLP, and LH (green and cyan in Figure 7). In this case, the most straightforward biological interpretation is the correspondence of the size of the sensory periphery (compound eye, in this case) with the size of the central targets; eyes of burrowing T. castaneum and the flying D. melanogaster differ in size considerably and so do the respective processing areas.

In other instances, the explanation for more subtle differences in structure is not as evident. In the present analysis, we found three cases where *T. castaneum* deviated clearly from the well-known *D. melanogaster* pattern, namely the trajectories of the loSL and obP fascicles, and the position of the SuEC commissure. It is likely that differences in lineage size and developmental timing (heterochrony) may explain some divergence between the species. For example, the exact timing of when two different fibre systems (such as the peduncle and the obP, or the medial lobe and the SuEC) meet could provide scenarios in which the position of the systems relative to each other diverges. If for example obP fibres (CP lineages) meet the peduncle when it is formed by only few axons (as in *D. melanogaster*; VH, unpublished) they may all be allowed, when following a straight course, to cross over the dorsal surface of the peduncle; the numerical relationship between CP fibres and peduncle fibres at the time at which they meet may be different in *T. castaneum*. Of course, other more “profound” differences in the expression pattern of attractive or repulsive surface molecules may also play a role in modifying the exact path of the tract. It is clear that careful developmental studies that examine intermediate developmental stages need to be conducted to elucidate the mechanisms for the divergence beyond our speculations.

The evolutionary connection between shape differences and their functional consequences seems to be more difficult to understand. Despite efforts in vertebrate brain evolution to identify such links (Eliason et al., 2021; Sansalone et al., 2020), corresponding efforts in arthropod evolution are rare. We assume that selection acts firstly on dimensions such as cell number, proliferation rate, as well as changes, truncation, or multiplication of connections. These dimensions readily explain differences in size but have a more complex influence on shape. Shape differences might be secondary consequences of modifications in size which in itself might be consequential of these developmental dimensions, such that shape differences might be a lag and not a lead effect. Therefore, while we acknowledge the large amount of shape differences, we refrain from interpreting these at this point.

### Potential conservation and divergence in early neural development

A small but growing number of studies in insects outside *D. melanogaster* indicate that the process of neuroblast formation and the expression pattern of transcription factors in brain neuroblasts may be strongly conserved among many insect species (Biffar & Stollewerk, 2014; Birkan et al., 2011; Boyan et al., 2017; Farnworth et al., 2020; Koniszewski et al., 2016; Truman & Ball, 1998), and, as a result, we may speak of genetically as well as functionally homologous lineages. This is most certainly the case for the lineages generating the central complex (DM1-4, DALcl1/2, DALv2) and the mushroom body, but most likely extends to lineages in general. It should be noted that the similarities in axon tract systems established here for *T. castaneum* and *D. melanogaster*, even without the availability of any specific developmental or gene expression data for the neurons in question, strongly point to the expression of similar genetic programs controlling neural morphogenesis. We observe, for example, that a subset of BAla lineages contacts the antennal lobe at its ventrolateral corner, before projecting medially into the mALT. This implies that during development, these BAla neurons in *T. castaneum* and *D. melanogaster* not only “recognize” the same specific (maybe even transient) target structures among other neurons that tell them where to enter the neuropil, but that the initial branching event (dendrites leaving the fibre in <u>anterior</u> direction, to produce an antennal lobe located <u>anterior</u> of the point of entry of the bundle) is also the same in both species, and thereby, presumably, controlled by similar genetic regulatory pathways unfolding in the growing fibres.

The best studied mechanism by which diversity in insect brain structure is achieved is neuroblast proliferation (Farris & Van Dyke, 2015; Koniszewski et al., 2016; Lee, 2017; Pop et al., 2020; Yaghmaeian Salmani & Thor, 2020). Neuroblasts may enter or exit their phase of asymmetric division at different time points. In addition, neuroblasts may duplicate (e.g., 2 vs 4 mushroom body lineages in *T. castaneum* and *D. melanogaster*; the paired lineages DALcm1/2, DALcl1/2). Apoptotic cell death may truncate lineages or eliminate entire hemilineages (Kumar et al., 2009; Pop et al., 2020; Prieto-Godino et al., 2020; Truman et al., 2010). In addition, we dare to speculate that there might be a mechanism to convert a neuroblast from type 1 to type 2, generating a large “clan” of intermediate progenitors from one ancestral type 2 neuroblast. The description of hundreds of proliferating neural progenitors (originally described as neuroblasts) in the large bee mushroom body (Farris et al., 2004) could represent a case of such “type 1>type2” transformation.

In the present study, we did not systematically quantify neuroblast proliferation. However, some salient features are worth pointing out because they may shed light on different strategies in neurogenesis followed by different members of the holometabolan clade. First, with the exception of the mushroom body neuroblasts, we found no other morphologically recognizable neuroblasts during the larval period. Nevertheless, the brain grows substantially during the larval stages. This lets us surmise that the secondary phase of neuroblast proliferation in this species may proceed in a less time-restricted way than in *D. melanogaster*. Rather than all neuroblasts entering the secondary proliferation phase at around the same time, and remaining active at the same time, larval neuroblasts in *T. castaneum* may be active for shorter periods of time and in a less coordinated way throughout larval life. It is also possible that some brain neuroblasts of *T. castaneum* lack a secondary phase of proliferation entirely, similar to the abdominal and (many of the) gnathal neuroblasts in *D. melanogaster* (Urbach et al., 2016). Furthermore, dividing neuroblasts might be difficult to see if they were smaller and remained located in midst of their progeny, embedded in the brain cortex rather than at the periphery. This would account for the position of (presumed) secondary neurons at any cortical depth. A systematic study of proliferation patterns will be required to distinguish between these possibilities.

Second, we observed that the synapse-free lacunae, a characteristic feature of the late larval *D. melanogaster* brain caused by the massive bundles of undifferentiated secondary axons and their filopodia, are not detectable in *T. castaneum*. This finding can be interpreted in either of two ways. (1) Secondary neurons in *T. castaneum* may not enter a phase of differentiative arrest as in flies but immediately differentiate and integrate in the existing circuitry, as was observed for neurons of the central complex (Farnworth et al., 2020). (2) Secondary neurons are fewer in number and are more scattered throughout the brain (which would coincide with the scarcity of neuroblast-like cells mentioned above). As a result, even if these scattered cells remained undifferentiated during the larval period, their presence would not lead to large coherent volumes of tissue that lack synapses, as we see in *D. melanogaster*.

It might be conceivable, however speculative, that these two factors – a halt of differentiation, and a time-coordinated and anatomically ordered location of neuroblasts – are connected in *D. melanogaster* development. Maybe, a distal location of the neuroblast allows for tighter and more invariant packaging of daughter cells which in turn allows for a rapid proliferation and a prefiguring *before* differentiation occurs (like in the CX, Andrade et al., 2019) as well as to reduce metabolic costs. With it, the larval phase can be shortened while still having prefigured the brain’s cells for a rapid differentiation in the pupal phase. Experimental verification is needed, possibly comparatively, to understand whether daughter cell density is a function of neuroblast location, and whether this is connected to developmental speed and a condensed larval period such as in *D. melanogaster*. Interestingly, the very emergence of holometabolous life cycle required a solution to the problem of completing the brain postembryonically. Our observation indicates that there is evolutionary flexibility in solving that problem, calling for dedicated studies regarding the developmental underpinnings.

### Heterochronies in brain development might be linked to different manifestations of holometaboly

Differences in the timing of neural proliferation and differentiation as discussed above are examples of a developmental-evolutionary mechanism called heterochrony. The impact of heterochrony on insect brain development has been previously shown for the central complex, which is present in the larva of *T. castaneum* and many other insect taxa but absent in larvae of *D. melanogaster* and other taxa (Farnworth et al., 2020; Koniszewski et al., 2016; Panov, 1959; Wegerhoff & Breidbach, 1992). The more continuous neural proliferation and/or differentiation found in *T. castaneum* central complex development may reflect similar behaviour of most if not all regions of the *T. castaneum* brain and fits with out general observations with respect to neuroblast proliferation and differentiation differences.

We wondered whether the timing differences in neural proliferation and differentiation are correlated with, and possibly causally related to, other aspects of development of *T. castaneum* and *D. melanogaster*. The development of the bodywall and its complement of peripheral sense organs is likely coordinated with the development of the central nervous system. In *D. melanogaster*, a similar decoupling of proliferation and differentiation observed for brain development is observed in the bodywall. Fly larvae are acephalic, lacking typical head structures, and possess no legs. Sensory neurons associated with the larval body wall do not increase in number. At the same time, future adult tissues are prefigured inside the larval body as imaginal discs, whose undifferentiated, proliferating cells do not participate in larval body functions. This is quite similar to future adult brain structures, which remain non-functional but are visible in the form of lacunae. By contrast, *T. castaneum* has an eucephalic head with simple eyes, antennae and mouth parts, and thoracic segments with short walking legs, thus reflecting much more the ground plan of the ancestral holometabolous larva (Peters et al., 2014) and the larva being, therefore, much more similar to the adult. Interestingly, *T. castaneum* larval epidermis seems to be carried over to the adult (Snodgrass, 1954; Jindra M., pers. comm.), indicating a more continuous development, at least with respect to this aspect.

What might the evolutionary advantage be for developing a fast-developing acephalic, leg-less larva as found most prominently in *D. melanogaster* and other flies? Maddrell (2018) and Truman & Riddiford (1999) hypothesize that the shedding of cuticle during molting can occur more rapidly if the larva has a simple cylindrical shape. As a result, larval development can be sped up. Indeed, *D. melanogaster* borrows itself in the food source, which reduces the necessity for structures and behaviors serving predator evasion. In contrast, *T. castaneum* and most other holometabolan insects have a much more extended larval phase (in absolute and relative terms, see Farnworth et al., 2020; Sokoloff, 1972): 13 to 23 days in *T. castaneum* and 4-5 days in *D. melanogaster*. In other words, *T. castaneum*, which undergoes a prolonged larval development requiring more complex behaviour to avoid predators or search for an appropriate food source, needs larval legs, antennae, and corresponding brain structures. This potential connection between feeding style, complexity of larval body, length of development and coupling of neural proliferation and differentiation illustrates the urgent need to analyze development and behaviour in alternative model organisms, particularly because *D. melanogaster* seems derived not only with respect to larval brain development. For example, it would be fascinating to examine imaginal disc and brain development in *Apis mellifera*, a species with a highly simplified larva as well and rapid larval development with even less demands for CNS function. The emerging genome editing tools allow labelling of homologous cell groups throughout development in different organisms, such as genetic neural lineages (Farnworth et al., 2020; Koniszewski et al., 2016) such that the developmental heterochronies can be traced and studied with high precision. Combining specific genetic marking with tools to generate clones has the potential to distinguish between cells of embryonic, larval or pupal origin.

### A Tubulin-based atlas as a gateway to a developmental view of the insect brain

With this work we provide a unique dataset that can provide a framework for future studies. First, the dataset does include not only the adult brain but also two important stages of holometabolan life cycle: the hatchling and the last larval stage. In contrast to other brain atlases, which focused on the adult brain only, this approach provides insights into the developmental origins of adult brain differences. Second, we compare not only neuropils and brain areas but the macro-circuitry as well. This aspect increases the depth of comparison and allows specifically labelled neurons to be assigned to certain fascicles, tracts and lineages groups. Indeed, having identified tracts in the hatchling and assigning these to neural lineage groups offers a well-needed connection to developmental processes in the insect brain. Given, that the antibodies used in this study work in most species, our method can easily be transferred. Third and most importantly, this work opens the stage for an analysis of the cellular and genetic bases of brain evolution in that it allows for more robust homology assignments of neurons between organisms. The exact description of the divergences of homologous cell’s development within a conserved framework is the prerequisite to find the genetic regulation of such divergence.

*T. castaneum* is an excellent model system for this matter because it is the insect model system closest to *D. melanogaster* with respect to versatility of genetic and transgenic methods. Specifically, transgenic methods including the Gal4 system (Berghammer et al., 1999; Schinko et al., 2010) can be used to establish reporters – based on this, use of a split Gal4 system should be possible. Genome editing via CRISPR/Cas9 is well established which opens exciting new possibilities. For instance, it has become possible to generate transgenic lines, where fluorescent proteins are driven in the patterns of homologous genes. In previous work, we have used a highly conserved transcription factor to label homologous genetic neural lineages, which represent a new unit of comparison previously not available (Farnworth et al., 2020; Koniszewski et al., 2016). The interpretation of such marked subsets of cells is greatly facilitated by this atlas and allows to focus on the identified differences with ease. The possibility of time- and cost-efficient RNAi screens in *T. castaneum* will facilitate the identification of the genetic control of development and evolutionary modification.

## Author contributions

MSF, GB and VH initiated the project. MSF performed the immunohistochemistry stainings, VH performed the analyses in TrakEM2. MSF and VH analysed and interpreted the data. GB, MSF and VH wrote the manuscript.

## Acknowledgements

We thank Prof. Christian Wegener for providing us with the anti-Synapsin antibody, and the Twitter Community for tips regarding immunohistochemistry improvements. MSF is supported by a Walter-Benjamin Fellowship from the Deutsche Forschungsgemeinschaft (FA1818/1-1). GB is supported by the Deutsche Forschungsgemeinschaft (BU1443/17-1) and VH by a National Institutes of Health grant (R01 NS054814-14).

## Competing interests

The authors declare that there are no competing interests.

## Notes

https://insectbraindb.org/

### Competing Interest Statement

The authors have declared no competing interest.

